# Autophagy acts as a spatial organizer of cell-type-specific plant immunity

**DOI:** 10.64898/2026.04.06.716646

**Authors:** Shanshuo Zhu, Manuel González-Fuente, Ophélie Léger, Gautier Langin, Ke Xu, Nazife Aydin, Nico Schulz, Pavel Solansky, Tom Denyer, Alessia Del Chiaro, Xinchao Wang, Qiangnan Feng, Yasin Dagdas, Marja Timmermans, Suayib Üstün

**Affiliations:** Faculty of Biology and Biotechnology, Ruhr-University Bochum; Bochum, 44780, Germany; Key Laboratory of Synthetic Biology of Ministry of Agriculture and Rural Affairs, Tobacco Research Institute, Chinese Academy of Agricultural Sciences, Qingdao 266101, China; Center for Plant Molecular Biology (ZMBP), University of Tübingen; Tübingen, 72076, Germany; Gregor Mendel Institute (GMI), Austrian Academy of Sciences, Vienna BioCenter; Vienna, 1030, Austria; State Key Laboratory of Wheat Improvement, College of Life Sciences, Shandong Agricultural University, Tai’an, China; Heidelberg University, Centre for Organismal Studies (COS), 69120 Heidelberg, Germany; Cluster of Excellence GreenRobust, Heidelberg University, 69120 Heidelberg, Germany

## Abstract

Effective plant immunity requires precise spatial coordination of immune responses across cell-types to restrict pathogens while preventing tissue damage, yet the intracellular pathways that organize this spatial architecture remain unknown. Here we show that autophagy serves as a spatial coordinator of immunity, partitioning and calibrating immune strategies across tissues during *Pseudomonas syringae* infection in *Arabidopsis thaliana*. Combining single-cell transcriptomics, cell-type-specific complementation, and live-cell imaging, we uncover distinct and opposing roles of autophagy across tissues. In guard cells, autophagy promotes pathogen-induced stomatal reopening by suppressing abscisic acid (ABA) signalling in part via autophagic turnover of the guard cell ABA receptor PYL4. In contrast, in mesophyll cells, autophagy restricts immune activation while simultaneously enabling effective immune execution: its loss enhances EDS1-PAD4-ADR1 pathway expression yet compromises PTI outputs. Mechanistically, bacterial infection triggers autophagic EDS1 turnover, supporting a model in which controlled EDS1 recycling facilitates HR elicitation. Cell-type-specific complementation reveals that PTI competence may involve coordinated autophagy across tissues, whereas mesophyll-autonomous autophagy is partially sufficient for HR execution. Together, these findings establish a framework in which spatial control of proteostasis orchestrates multicellular immune coordination, with broad implications for understanding how conserved degradation pathways regulate layered immunity across organisms.

## Main

Plant immune defences must be robust enough to stop pathogen proliferation yet carefully controlled to avoid overactivation, requiring an intricate and multilayered immune system. In the first layer, pathogen-associated molecular patterns (PAMPs) are perceived by cell surface receptors to activate pattern-triggered immunity (PTI), leading to transcriptional reprogramming and stomatal closure to limit bacterial entry into the apoplast^1^. Adapted pathogens overcome PTI by delivering effector proteins into host cells that can subvert this first layer of defence and promote infection^2^. In turn, plants evolved the ability to monitor some of these effectors by intracellular immune receptors that activate effector-triggered immunity (ETI)^1^. ETI amplifies PTI outputs and is frequently accompanied by the hypersensitive response (HR), a localized cell death that restricts pathogen growth at the infection site^3^. Accumulating evidence indicates that PTI and ETI are not independent immune layers but instead mutually potentiate each other to generate more robust defence outputs^3–6^. However, the cellular pathways that coordinate this immune potentiation remain poorly understood.

The complex molecular mechanisms during plant immunity require a high degree of proteomic flexibility. As such, cellular proteostasis pathways such as autophagy have emerged as important regulators of plant–microbe interactions^7,8^. Autophagy is a conserved degradation pathway that delivers cytoplasmic content to the vacuole for degradation^9^. Autophagy has also emerged as a central player in plant-microbe interactions, where it can be exploited by pathogens and is involved in the induction of pathogen-associated cell death^10–12^. However, at the same time, canonical PTI outputs are not altered in autophagy mutants^13^, although these early studies did not account for the cell-type heterogeneity during plant immune reactions^14^.

One explanation for this ambiguity may lie in the spatial organization of plant defence responses. During infections, bacterial pathogens such as *Pseudomonas syringae* pv. *tomato* DC3000 (*Pst*) enter leaves predominantly through stomata, making guard cells the first barrier encountered by the pathogen^15^. While stomatal closure restricts bacterial invasion, successful pathogens can actively induce stomatal reopening to enter the host^16^. At later stages of infection, mesophyll cells mount immune responses that aim at preventing bacterial proliferation from within^17^. These observations suggest that immune responses are spatially organized across tissues and cell types during infection. However, how intracellular pathways coordinate these distinct immune programs remains largely unknown.

Recent advances in single-cell transcriptomics have revealed extensive heterogeneity in plant immune responses, with individual cell populations adopting distinct transcriptional states^17–19^. As such, plant immunity is spatially organized at cellular resolution rather than uniformly distributed across tissues and cell types. However, most single-cell analyses provide limited insight into the cellular pathways that could control spatial organization and thereby regulate these heterogeneous immune states. Therefore, whether and how intracellular regulatory mechanisms, including degradation pathways, shape cell-type-specific immune responses during infection remains elusive.

Here, we combine single-cell RNA sequencing, cell-type-specific genetic complementation, and live-cell imaging to investigate the role of autophagy during *Pseudomonas syringae* infection in *Arabidopsis thaliana* at the cellular resolution. Our analyses reveal that autophagy performs distinct functions across different tissues during bacterial infection. In guard cells, autophagy promotes pathogen-induced stomatal reopening through selective degradation of the ABA receptor PYR1-LIKE PROTEIN 4 (PYL4). In contrast, autophagy deficiency primes immune responses in mesophyll cells, resulting in constitutive activation of the immunity-related ENHANCED DISEASE SUSCEPTIBILITY 1 (EDS1)-PHYTOALEXIN DEFICIENT 4 (PAD4)-ACTIVATED DISEASE RESISTANCE 1 (ADR1) pathway. However, despite elevated EDS1-dependent signalling, canonical PTI outputs and ETI execution remain compromised. Cell-type-specific complementation reveals that PTI competence may involve coordinated autophagy across tissues, whereas mesophyll-autonomous autophagy is partially sufficient for HR execution. Mechanistically, we show that bacterial infection itself triggers the autophagic degradation of EDS1, and that this controlled turnover is essential for productive immune execution. Together, our findings identify autophagy as a spatial organizer that partitions immune strategies across plant tissues during infection and contributes to both pathogen-mediated stomata reopening and EDS1-dependent convergence of cell-surface and intracellular immune perception.

## Results

### Autophagy acts in a cell-type-specific manner

Previous studies reported contrasting roles of autophagy in plant–microbe interactions, including functions that promote susceptibility^20–23^, likely because pathogen infection strategies and cell type–specific immune responses were not considered, features only recently resolved by single-cell transcriptomics^14^. Since *Pst* enters through stomata to then proliferate at later stages of infection in the mesophyll, we asked whether autophagy spatially organizes immune responses at this first line of defence.

Autophagy-deficient *atg5* and nonuple *atg8* mutants showed reduced bacterial growth upon dip inoculation (Fig. 1a), a phenotype compromised by syringe infiltration that bypasses stomatal entry (Extended Data Fig. 1a). Conversely, autophagy deficiency enhanced disease-related chlorosis upon infiltration, indicating a possible distinct function in mesophyll cells (Extended Data Fig. 1b). These results suggest cell-autonomous roles of autophagy in both guard and mesophyll cells. To further investigate this, we used a transcriptional reporter, *pATG8e::nls-3xGFP*, and observed rapid autophagy induction across cell types at 6 hours post infection (hpi), followed by sustained activation in mesophyll cells but attenuation in guard cells at 48 hpi (Fig. 1b). Together with the guard cell–specific expression and infection-induced upregulation of ATG8c^24^(Extended Data Fig. 1c), these data indicate that *Pst* infection elicits distinct autophagy dynamics across cell types, suggesting that autophagy may contribute to the spatial regulation of immune responses.

**Fig. 1:**
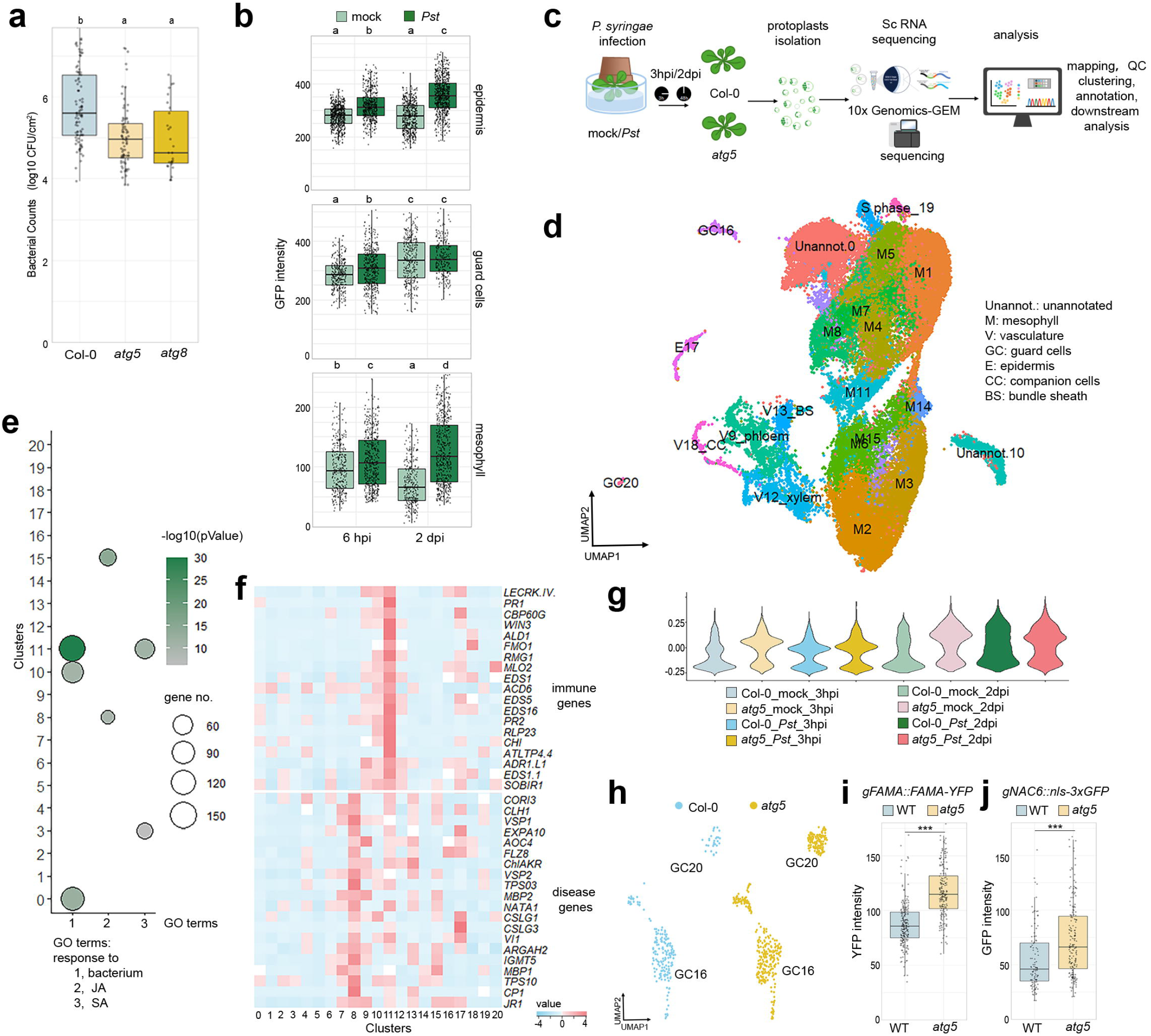
Autophagy displays cell-type-specific responses upon infection. **a**, Bacterial population in 5-week-old *A. thaliana* leaves of wild type Col-0 and *atg5, atg8* plants dip-inoculated with 10^8^ CFUs/mL *Pst*. Data from more than four independent experiments. Different letters indicate statistical groups determined using one-way ANOVA, followed by a Tukey HSD post-hoc test (adjusted p < 0.05). **b**, GFP fluorescence intensity detected from epidermis, guard and mesophyll cells of 5-week-old *A. thaliana* leaves expressing *pATG8e::nls-3xGFP* at 6 hours and 24 hours upon dip-inoculation with 10^8^ CFUs/mL *Pst* or 10 mM MgCl_2_ mock. For each cell type, letters indicate statistically significant differences determined by two-way ANOVA, followed by a Tukey HSD post-hoc test (adjusted p < 0.05). Experiments were performed three times with similar results. **c**, Schematic of the scRNA approach. **d**, Two-dimensional embedding of cells from all 8 samples by uniform manifold approximation and projection (UMAP) based on the transcriptomic data. Cells are coloured according to clusters. Cell types were annotated on the basis of marker gene expression. **e**, GO enrichment analysis for marker genes of each cluster (from Fig. S1e). GO terms related to defence are shown. p values from Fisher’s exact test are shown. Size of shape stands for the number of marker genes associated with the GO term. **f**, Heat map showing relative expression of defence-and disease-related genes across cell populations in different clusters. **g**, Violin plots showing the distribution of the Pst-responsive score across 8 samples. The shape illustrates the density of the score distribution for each sample. **h**, UMAP visualization of guard cells, with cells colored by genotype. **i**,**j**, Fluorescence signal intensity from guard cells in leaves or cotyledons of 10-day-old *A. thaliana* expressing *pFAMA::gFAMA-YFP* (I) and *pNAC6::nls-3xGFP* (J) in wildtype and *atg5* background. Statistical differences between genotypes are assessed withStudent’s t-test (\*\*\**p* < 0.001).

To resolve the cell type–specific role of autophagy in plant-bacteria interactions, we performed single-cell RNA sequencing (scRNA-seq) of Col-0 and *atg5* at early (3 hpi) and late (2 dpi) stages of *Pst* infection (Fig. 1c). Across eight samples, we obtained 54,039 high-quality cells and resolved distinct cell types (Fig. 1d). Several clusters were selectively enriched in the mutant or at specific infection stages (Extended Data Fig. 1d), indicating that clustering captured discrete cell states shaped by pathogen challenge and genotype. Marker gene identification (Table S1) followed by Gene Ontology (GO) enrichment revealed that mesophyll cluster M8 was enriched for disease-related genes associated JA responses (Fig. 1e; Extended Data Fig. 1e), whereas M11 was associated with response to bacteria, SA genes (Fig. 1f). Both clusters were predominantly observed at 2 dpi after *Pst* treatment (Extended Data Fig. 1f), indicating that these cells were actively responding to bacteria. This was further supported by elevated expression of canonical immunity genes in M11 and disease-associated genes in M8 (Fig. 1f). Integration with a published scRNA-seq dataset of *Pst*-infected tissue showed that M11 co-clustered with previously defined immunity-associated states (M1–M3), whereas M8 aligned with susceptible states (M4–M5) (Extended Data Fig. 1g). Together, these data define M11 as an immunity-associated and M8 as a disease-associated cell state. We next computed a signature score (“*Pst*-responsive score”) to quantify the impact of infection at single-cell resolution, using module scores derived from genes identified as differentially expressed in our bulk RNA-seq dataset (Table S2). Cells with high *Pst*-responsive scores were strongly enriched in infected samples at both 3 hpi and 2 dpi (Fig. 1g). Notably, a subset of cells in mock-treated *atg5* already displayed elevated scores, indicating a constitutively pre-activated immune state in the absence of pathogen (Fig. 1g). To further characterize this pre-activation, we identified differentially expressed genes (DEGs) upregulated in *atg5* compared to Col-0 across cell types under mock conditions (Extended Data Fig. 1h; Table S3). Shared DEGs across all cell types were strongly enriched for responses to bacteria. In contrast, cell type–specific DEGs revealed distinct signatures: mesophyll cells were enriched for salicylic acid (SA)-related processes, guard cells for ABA responses, and epidermal cells for ethylene-and jasmonic acid (JA)-mediated signalling (Extended Data Fig. 1h), highlighting the role of autophagy in cell type– specific defence strategies.

Consistent with the emergence of cell type–specific defence strategies, we next focused on guard cells to corroborate our findings that autophagy regulates the contribution of stomata to plant immunity. We resolved two guard cell clusters defined by established developmental markers (Extended Data Fig. 2a). The G16 cluster comprised cells from both Col-0 and *atg5* and aligned with guard cells from the integrated reference dataset (Extended Data Fig. 1g), whereas GC20 was predominantly composed of *atg5*-derived cells (Fig. 1h; Extended Data Fig. 1d), suggesting a mutant-specific guard cell state. To assess whether this shift reflects altered guard cell development, we introduced *pSCAP1::nls-mTurquoise2* and *pFAMA::gFAMA-YFP* reporters into the *atg5* background^25^. Both markers showed increased expression in *atg5* (Fig. 1i; Extended Data Fig. 2b), consistent with altered guard cell developmental regulation in the mutant and potentially explaining the increased guard cell abundance in the mutant. Strikingly, genes enriched in GC20 relative to G16 were associated with stress-responsive pathways, including hypoxia, bacterial defence, and ABA signalling (Extended Data Fig. 2c), indicating a defence-activated guard cell state. Focusing on GC20-specific signatures, we identified *atg5*-enriched genes with high expression in this cluster (Table S4). Among these, members of the SERINE-RICH ENDOGENOUS PEPTIDES (SCOOP) family, known to trigger reactive oxygen species (ROS) production, Ca^2+^ signalling, and immunity^26–28^, were prominently represented, with *SCOOP14* highly expressed in GC20, E17, and the immune-associated mesophyll cluster M11 (Extended Data Fig. 2d, e). Reporter analysis (*pSCOOP14::nls-3xGFP*) confirmed elevated expression in *atg5* guard cells (Extended Data Fig. 2h,i). In addition, the NAC transcription factor *NAC6/ORE1/NAC092*, previously linked to stress responses and age-dependent cell death^29,30^ showed strong and specific expression in GC20 and was elevated in *atg5* (Extended Data Fig. 2f, g). This was validated using a *pNAC6::nls-3xGFP* reporter, which also revealed induction upon *Pst* infection (Fig. 1j; Extended Data Fig. 2j, k). Together, these data indicate that autophagy deficiency drives the emergence of a distinct, defence-primed guard cell state.

### Autophagy promotes stomatal opening during infection

Considering the contrasting bacterial growth phenotypes and the distinct guard cell-specific defence state we observed in autophagy deficient plants, we sought to investigate the expression of autophagy-related genes (*ATGs*) in guard cells. Utilizing our scRNA-seq data, we observed prominent expression of most *ATGs* in guard cells than in other cell types, supporting our findings that autophagy plays a specific role in guard cells (Fig 2a). Given that stomatal immunity is critical for limiting *Pst* invasion, we next assessed stomatal aperture in *atg5* during infection. Similar to Col-0, stomata in *atg5* closed at 1 hpi; however, re-opening was impaired at 3 hpi, a critical window for bacterial invasion^31^ (Fig. 2b; Extended Data Fig. 3a).

**Fig. 2:**
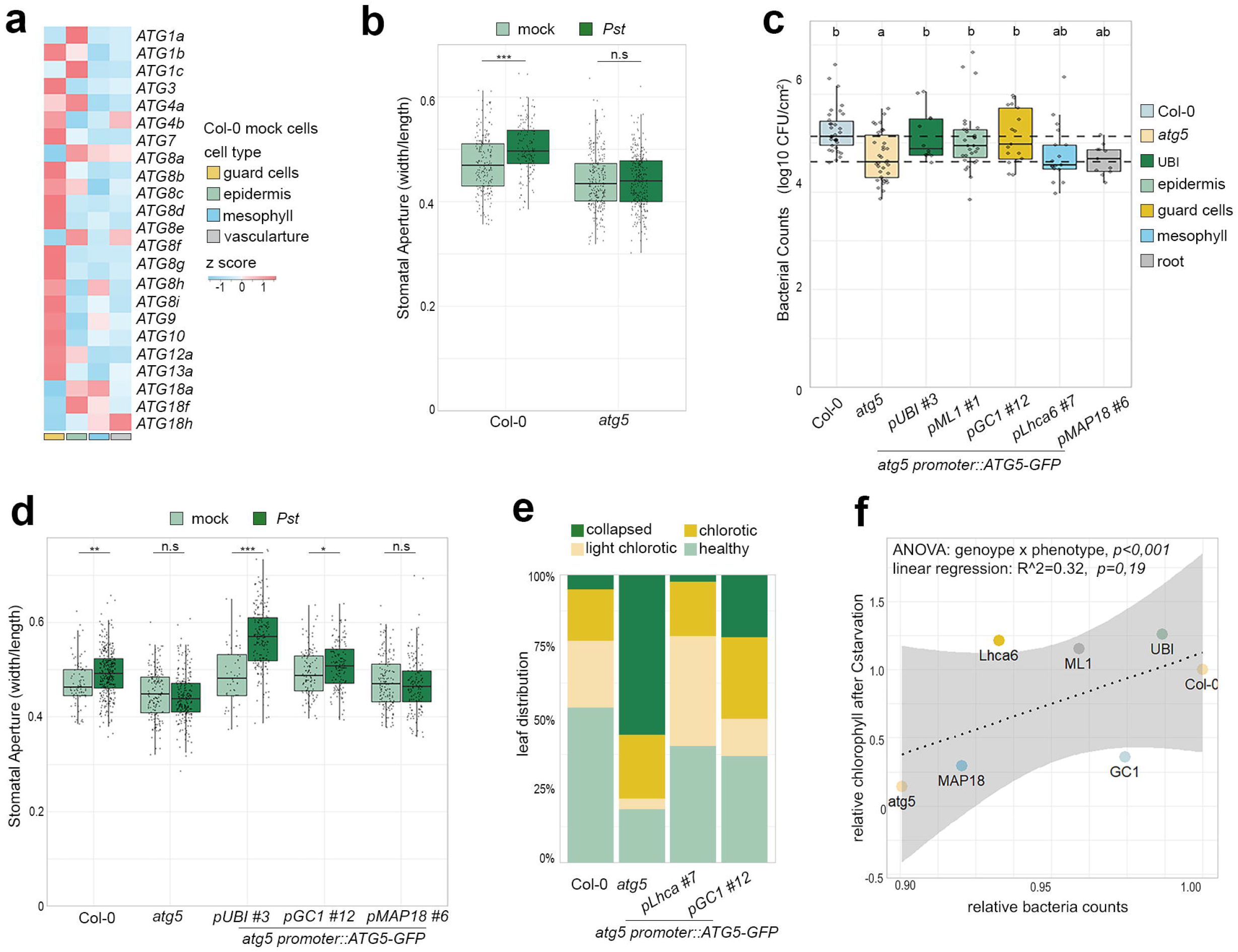
Autophagy promotes stomatal opening during infection. **a**, Heat map showing relative expression of autophagy-related genes across four major cell types in Col-0 mock samples. **b**, Stomatal apertures of Col-0 and *atg5*plants dip-inoculated with 10^8^ CFUs/mL *Pst* or mock at 3 hours. Statistical differences between treatment within each genotype are determined by two-way ANOVA followed by Tukey HSD post-hoc tests (adjusted \*\*\**p* < 0.001, n.s, not significant). This experiment was repeated at least three times with similar results. **c**, Bacterial growth in Col-0, *atg5* and *atg5* complementation lines carrying tissue-specific promoters after dip-inoculation with 10^8^ CFUs/mL *Pst*. Dashed horizontal lines indicate the medians of Col-0 and *atg5* across all experiments. Different letters indicate statistical groups determined using one-way ANOVA with experiment as a random effect, followed by a Tukey HSD post-hoc test (adjusted *p* < 0.05). Data from six independent experiments. **d**, Stomatal apertures of 5-week-old *A. thaliana* Col-0, *atg5* and *atg5* complementation lines carrying tissue-specific promoters. Leaves were taken and fixed at 3 hours after inoculation with 10^8^ CFUs/mL *Pst* or 10 mM MgCl_2_ mock. Statistical differences between treatment within each genotype are determined by two-way ANOVA followed by Tukey HSD post-hoc tests (adjusted \**p*<0.05, \*\**p* <0.01, \*\*\**p* < 0.001, n.s, not significant). This experiment was repeated three times with similar results. **e**, Leaf phenotypic analysis of Col-0, *atg5* and *atg5* complementation plants after syringe-infiltrated with 5 x 10^4^ CFUs/mL *Pst*. Stacked bar chart illustrates the distribution of leaf states (collapsed, chlorotic, light chlorotic, healthy) as percentages of total leaves per genotype. The *p* value was calculated using a chi-square test (χ^2^ = 27.785, df = 9, p-value = 0.001036). This experiment was repeated three times with similar results. **f**, Relationship between bacterial growth complementation and carbon starvation rescue for *atg5* complementing lines, together with Col-0 and *atg5*. Each point represents the relative value to wild type (x-axis, relative log CFU to Col-0; y-axis, relative chlorophyll retention to Col-0) for a given genotype, colored as indicated. The dashed line shows the linear regression fit across all data points (*R*^2^ = 0.32, *p* = 0.19). The *p*-value for the genotype-by-phenotype interaction (two-way ANOVA, Type III) is shown ().

In parallel, we identified a mesophyll cluster (M14) specifically enriched in 3 hpi–infected samples (Extended Data Fig. 1d). Marker gene analysis revealed enrichment for stress-related processes, including hypoxia, wounding, and ethylene signalling (Extended Data Fig. 3b), suggesting that M14 represents an early-responsive mesophyll population encountering invading bacteria. Notably, this cluster was less enriched in *atg5* at 3 hpi, consistent with impaired stomatal re-opening of autophagy deficient mutants limiting bacterial entry and consequently reducing early mesophyll engagement.

To further investigate the cell-type-specific role of autophagy, we introduced the coding sequence (CDS) of ATG5 driven by distinct cell-type promoters into the *atg5* mutant background (Extended Data Fig. 3c). We observed that the *pUBI::ATG5-GFP* (constitutive), *pML1::ATG5-GFP* (epidermis-specific) and *pGC1:: ATG5-GFP* (guard cell-specific) lines could rescue the enhanced resistance phenotype of *atg5*, whereas *pLhca6::ATG5-GFP* (mesophyll-specific) and *pMAP18*::*ATG5-GFP* (root-specific) lines behaved like the *atg5* mutant (Fig 2c, Extended Data Fig. 3d). This was also consistent with the stomatal aperture measurements, since we found that the expression of *pGC1::ATG5-GFP* and *pUBI::ATG5-GFP* in *atg5* restored the stomatal re-opening phenotypes, whereas the *pMAP18::ATG5-GFP atg5* complementation line maintained smaller stomatal apertures upon *Pst* infection similar to the *atg5* mutant (Fig. 2d). In contrast, only mesophyll-driven *atg5* complementation prevented enhanced tissue collapse and chlorosis after syringe infiltration while guard cell-specific complementation did not (Fig. 2e), indicating autophagy may partition defence responses between guard cells and mesophyll.

To assess whether this cell-type-specific response is distinct to infection or other physiological functions of autophagy, we monitored early senescence phenotypes for the same lines and subjected them to nutrient starvation. Our analysis revealed that *pUBI*::, *pML1*::and *pLhca6* driven *ATG5-GFP* expression fully complemented these phenotypes, whereas the *pGC1::* or *pMAP18*::*ATG5-GFP* did not or only partially complemented these phenotypes (Extended Data Fig. 3e-g). Linear regression analysis revealed a moderate, but non-significant, positive relationship between bacterial growth and carbon starvation complementation across promoters (Fig. 2f). However, the relative performance of individual promoters differed markedly between the two phenotypes (immune response and carbon starvation), indicating tissue-specific functional specialization (Fig. 2f). Promoters deviating from the regression line displayed preferential complementation effects: *pLhca6*, positioned above the line, conferred stronger rescue under carbon starvation, whereas *pGC1*, below the line, showed the opposite trend. Notably, *pGC1* restored bacterial growth to near wild-type levels but failed to rescue chlorophyll retention, suggesting that guard cell–specific autophagy contributes to plant– pathogen interactions in a manner that is uncoupled from bulk leaf autophagic activity, which seems to be majorly executed by mesophyll and epidermal cells.

### Autophagy fine-tunes ABA activity by recycling a guard cell-specific ABA receptor

Given the role of ABA in regulating stomatal closure during infection^16^ and the enrichment of ABA-related genes in the guard cell cluster GC20 in *atg5* (Extended Data Fig 1h), we investigated whether autophagy mediates stomatal re-opening in an ABA-dependent manner. We first assessed the role of reduced ABA signalling in *Pst*-induced stomatal re-opening and analysed mutants with constitutively elevated ABA levels or responses: *cyp707a-1* (defective in ABA catabolism), shown to be essential for stomatal opening and Pst invasion^32,33^, and *ahg3-2* (ABA-hypersensitive)^34,35^. In both mutants, *Pst*-triggered stomatal reopening at 3 hpi was abolished (Extended Data Fig. 4a). Consistently, exogenous ABA application prevented stomatal re-opening upon infection (Extended Data Fig. 4b). Next, we generated double mutants with the ABA-deficient *aba2-1*^*16*^ and the ABA signalling mutant *ost1-3*^*36*^and revealed that reduced ABA signalling could restore the re-opening defect in *atg5* (Fig. 3a). This was further supported by pharmacological inhibition of ABA biosynthesis, as fluridone treatment partially rescued the *atg5* stomatal phenotype (Extended Data Fig. 4c). Taken together, our data demonstrate that autophagy promotes stomatal re-opening by suppressing the activity of ABA pathways during infection.

**Fig. 3:**
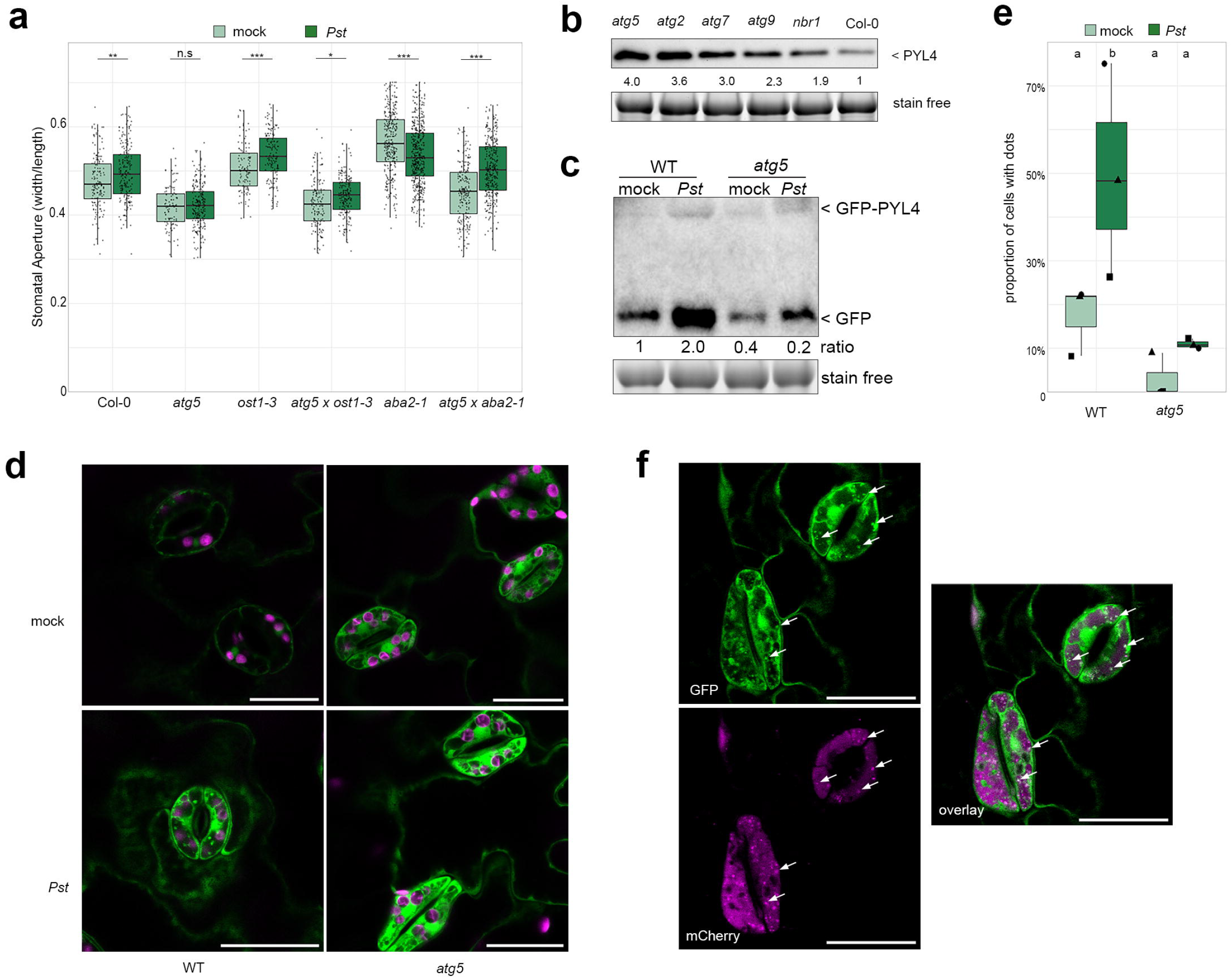
Autophagy fine-tunes ABA activity by recycling a guard cell-specific ABA receptor. **a**, Stomatal apertures of Col-0, *atg5, ost1-3, aba2-1* and double mutants *atg5 x ost1-3, atg5 x aba2-1* plants. 5-week-old *A. thaliana* leaves were taken and fixed at 3 hours after incubated with 10^8^ CFUs/mL *Pst* or 10 mM MgCl_2_ mock. Statistical differences between treatment within each genotype are determined by two-way ANOVA followed by Tukey HSD post-hoc tests (adjusted \**p*<0.05, \*\**p* <0.01, \*\*\**p* < 0.001, n.s, not significant). This experiment was repeated three times with similar results. **b**, Immunoblot analysis of PYL4 protein in 7-day-old *A. thaliana* seedlings of Col-0 and autophagy defective mutants. The large subunit of the Rubisco visible through Stain-Free imaging revelation serves as loading control. Numbers indicate the band intensity of PYL4 immunoblot normalized to each sample loading. The experiment was repeated three times with similar results. **c**, PYL4 degradation assay revealed by the ratio of free GFP to full-length GFP–PYL4. Immunoblot against GFP on crude extracts of 10-day-old *A. thaliana* seedlings GFP–PYL4 in wild type and *atg5* background, infected with 10^8^ CFUs/mL *Pst* or 10 mM MgCl_2_ mock for 5 hours. Numbers indicate GFP/GFP–PYL4 ratio. The experiment was repeated two times with similar results. **d**, Representative confocal microscopy images of 10-day-old *A. thaliana* wild type and *atg5* leaves expressing *GFP–PYL4*. Overnight 1 µm concanamycin A treated seedlings were incubated with 10^8^ CFUs/mL *Pst* or 10 mM MgCl_2_ mock for 2 hours. Scale bar = 25 μm. **e**, Quantification of the proportion of cells with dots for each condition from (F). Data represent three independent experiments, with each shape corresponding to an individual experiment. Statistically significant differences were determined using a binomial generalized linear mixed model with experiment as a random effect, followed by a Tukey HSD post-hoc test (adjusted p < 0.05). **f**, Confocal microscopy pictures of *A. thaliana* leaves expressing GFP-PYL4 with mcherry-ATG8e. Co-localization is visible in ATG8e-labelled autophagosomes (indicated by arrows). Scare bar = 25 μm.

Sustained ABA activity requires tight control of receptor abundance and localization, as ABA receptors (PYR/PYL/RCAR) undergo continuous ubiquitination-dependent trafficking and degradation via endosomal–vacuolar pathways^37^. Considering the role of autophagy as a vacuolar degradation pathway, we hypothesized it could recycle ABA receptors that have been previously described to be degraded in the vacuole^38,39^. To this end, we assessed ABA receptor protein levels in autophagy-defective mutants and found that PYR1, PYL3 and PYL4 accumulated in *atg2* and *atg5* (Fig. 3b, Extended Data Fig. 4d). Given that *PYL4* is predominantly expressed in guard cells according to our scRNA-seq data (Extended Data Fig. 4e), we focused on this receptor and confirmed its accumulation across additional autophagy mutants (Fig. 3b). Consistently, the reporter line *pPYL4::nls-3xGFP* showed dominant expression in guard cells compared to other epidermal cells (Extended Data Fig. 4f), with a modest increase at the transcript level in *atg5* (Extended Data Fig. 4g). To test whether PYL4 is targeted by autophagy during infection, we quantified the ratio of free GFP to full-length GFP–PYL4 as a proxy for degradation. Upon *Pst* infection, free GFP accumulated, indicating enhanced PYL4 turnover (Fig. 3e). In *atg5*, basal degradation was reduced and *Pst*-induced turnover was impaired, as reflected by a lower free GFP/GFP–PYL4 ratio under both mock and infection conditions (Fig. 3c). In accordance with this, *Pst* induced the formation of punctate GFP–PYL4 structures in guard cells, co-localizing with mCherry-ATG8E (Fig. 3d, f), which accumulated in the vacuole upon concanamycin A (ConA) treatment, indicating autophagic delivery. In contrast, these structures were absent in *atg5* and did not further accumulate after *Pst* infection under concanamycin A treatment (Fig. 3d and e), supporting the notion that PYL4, and possibly additional ABA receptors, are degraded by autophagy, thereby regulating ABA responsiveness and facilitating stomatal re-opening during bacterial infection.

PYL4 has been reported to undergo vacuolar degradation via the late endosome pathway, with the ESCRT-III–associated protein ALIX acting as an adaptor^38^. Since ALIX also resides in a complex with ATG8 during salt stress^40^, that can rapidly increase ABA levels, we hypothesized that ALIX may also link PYL4 to the autophagy machinery. In line with this, we observed that ALIX interacts with ATG8 in co-immunoprecipitation assays and co-localizes with ATG8-labelled structures (Extended Data Fig. 4h, i), supporting a role at the interface of endosomal trafficking and autophagy. To test the functional relevance of this, we examined the response of the *alix* mutant to *Pst* infection. Strikingly, *Pst*-induced stomatal reopening was abolished in *alix* (Extended Data Fig. 4j), phenocopying the defect observed in autophagy mutants. Together, these results support a working model in which autophagy contributes to PYL4 turnover, likely acting through the ALIX-ATG8 node, thereby enabling stomatal re-opening during *Pst* infection.

### Autophagy alters immune gene response in the mesophyll

Together, our findings so far establish a guard cell–specific role of autophagy in controlling pathogen entry. When this barrier is bypassed, however, autophagy-deficient plants display enhanced chlorosis and less bacterial growth (Extended Data Fig 1b), indicating that autophagy deficiency confers a robust defense strategy that extends beyond the guard cell, likely within mesophyll tissue. Consistent with this idea, our single-cell analysis uncovered specific autophagy-dependent immune states across cell types, including defence- and disease-associated mesophyll populations (Fig 1f). To define the mesophyll-specific defence mechanism, we focused on the immunity-associated cluster M11. Notably, a population of cells with high *Pst*-responsive scores was already present in mock-treated *atg5* but practically absent in mock-treated Col-0 (Fig. 4a), indicating an *atg5*-specific pre-activated immune state. Detailed analysis of these cells revealed a set of genes associated with enhanced immunity in *atg5* (Extended Data Fig. 5a, b and Table S5). This included well-characterized defence genes such as the SA-responsive marker (*PR1 and PR2*)^41^, ER-resident *SDF2* being required for PTI signalling^42^, key components of the signalling module linking PTI and ETI, the EDS1-PAD4-ADR1 node^5^, as well as *PEN3*, previously implicated in bacterial resistance^43^ (Fig. 4b and c). Using reporter lines, we confirmed increased expression of *EDS1, PEN3* and *SDF2* in the autophagy-deficient mutant and after immune activation in wild type via syringe-inoculation of *Pst ΔhrcC* (Fig. 4d, e; Extended Data Fig. 5c–f), which ensures more uniform immune activation and controls for *atg5*-dependent differences in bacterial entry. At cellular resolution, *SDF2, PR2* and *ADR1-L1* were little or rarely detected in mesophyll cells under mock conditions but were induced upon *Pst ΔhrcC* infection, with *atg5* displaying a higher proportion of mesophyll cells expressing these genes even in the absence of infection (Extended Data Fig. 5e-h). More broadly, most immune-related genes that were largely absent from mesophyll cells under mock conditions were instead enriched in vascular, guard, and epidermal cells (Extended Data Fig. 5i). Specifically, *SDF2* was highly expressed in guard and vascular cells, whereas *EDS1* was pre-dominantly detected in epidermal and vascular cells (Extended Data Fig. 5j, k). Similar expression patterns of other immune components in the vasculature have been described previously^19,44^. Together, these findings indicate that autophagy restricts the spatial deployment of immune signalling, and that its impairment leads to enhanced defence gene expression in mesophyll cells, establishing a pre-activated, cell type–specific immune state that enhances bacterial resistance.

**Fig. 4:**
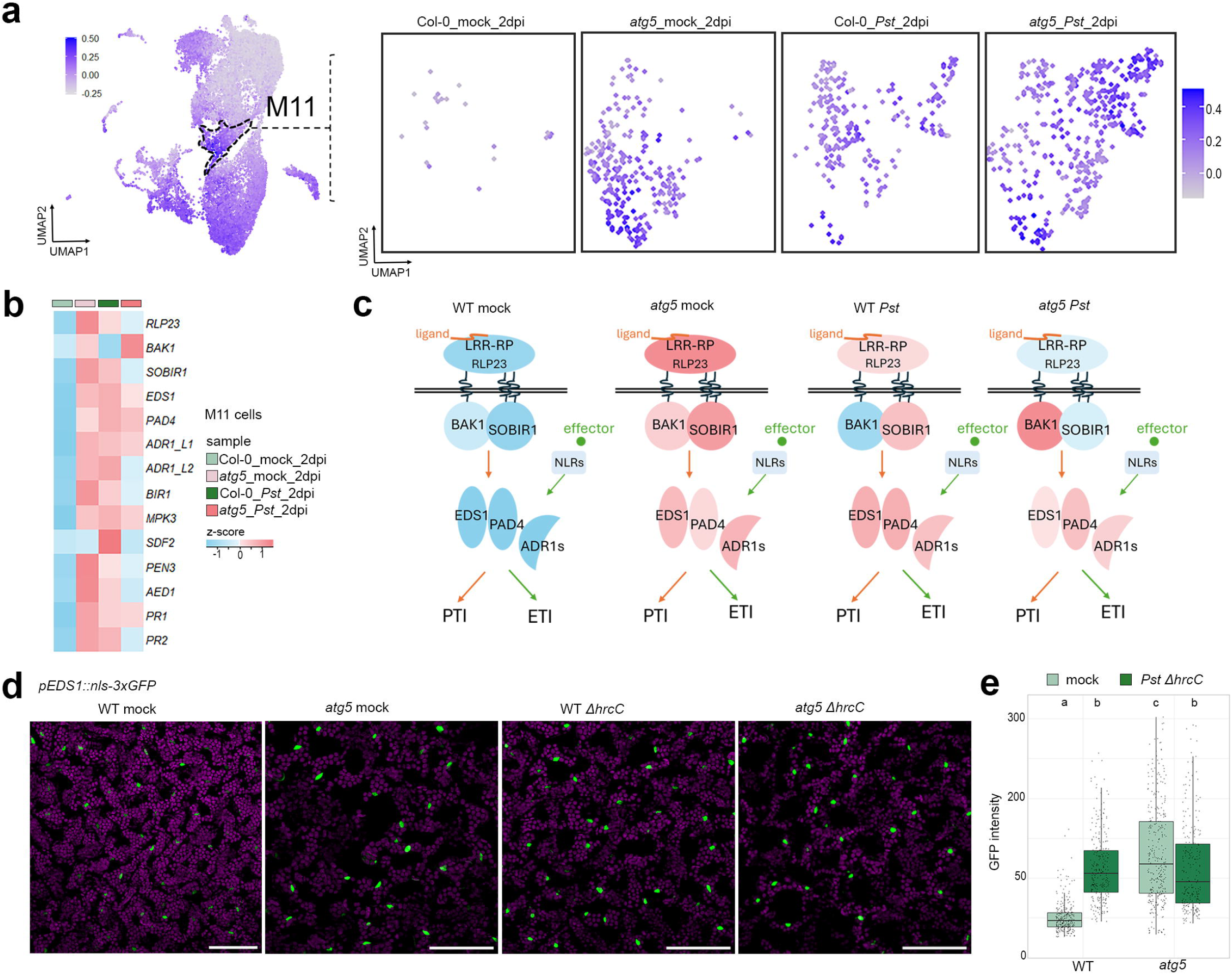
Autophagy alters immune gene expression in the mesophyll. **a**, UMAP plot visualizing Pst_responsive score across M11 cluster cells. Left panel: dashed line outlines the M11 immune clusters. Right panel: M11 UMAP for the four 2dpi samples: Col-0_mock _2 dpi, *atg5*_mock_2 dpi, Col-0_*Pst*_2 dpi, and *atg5*_*Pst*_2 dpi. **b**, heatmap showing the relative expression of selected *atg5*-immune-specific candidates in M11 cluster. **c**, The EDS1-PAD4-ADR signaling node as a convergence point of PTI and ETI, displayed the same across four experimental conditions. Schematic adapted from (Pruitt et al. 2021) to illustrate the immune signaling pathways. Protein colors correspond to the relative expression levels of indicated genes quantified in the heatmap (Fig. 4b). **d**, Confocal microscopy images of wild type and *atg5* plants expressing *pEDS1::nls-3xGFP*, syringe-infiltrated with 5×10^7^ CFUs/mL *Pst ΔhrcC* or or 10 mM MgCl_2_ mock. Leaves were imaged at 1 dpi and fluorescent images are maximal intensity projections of Z-stacks. Bar scale represents 100 μm. **e**, quantification GFP intensity of Fig.4D. Different letters indicate statistically significant differences, determined by two-way ANOVA followed by Tukey HSD post-hoc tests (adjusted p < 0.05). This experiment was repeated three times with similar results.

To further resolve cell type–specific immune signatures, we analyzed the disease-associated mesophyll cluster M8. This cluster was sparsely represented across mock conditions but became enriched upon infection. Notably, *Pst* challenge led to a higher proportion of M8 cells with elevated *Pst*-responsive scores in Col-0 compared to *atg5* (Extended Data Fig. 6a), indicating a reduced activation of this disease-associated state in the autophagy-deficient background. Transcriptional profiling of M8 cells between Col-0 and *atg5* revealed attenuated expression of JA-responsive genes in *atg5*, including *VSP* family genes^45^ (Extended Data Fig. 6b–d, and Table S6). Given the well-established role of JA signalling as a susceptibility hub in plant-bacteria interaction^46^, this suggests that autophagy promotes a JA-associated disease state within mesophyll cells. To validate these findings *in vivo*, we employed the reporter line *pVSP1::nls-3xGFP*. Following 4 days of *Pst* infection, a higher number of GFP-positive nuclei was observed in wild-type plants compared to *atg5*, whereas reporter activity was absent under mock conditions (Extended Data Fig. Fig. 6e, f). Together, these results demonstrate that autophagy is required for susceptibility and the proper establishment of the M8 disease-associated mesophyll state during bacterial infection.

### Autophagy plays a role in mesophyll-specific immune potentiation

Our reporter analysis of defence genes indicates activation in mesophyll cells independent of stomatal entry, supporting a mesophyll-specific role of autophagy during plant immunity. Among the defence genes enriched in M11, the pre-activated mesophyll population in *atg5*, components of the EDS1–PAD4–ADR1 signalling module were particularly prominent (Fig. 4c), a central node integrating cell surface and intracellular immune pathways^5^. Given its established role in PTI–ETI convergence, we asked whether autophagy modulates immune potentiation through this pathway. Consistent with our scRNA-seq expression data and reporter lines, we observed induced protein levels of EDS1 and MPK3 upon bacterial infection that was further elevated in autophagy deficient plants already under mock conditions (Fig. 5a). Further analysis of the protein abundance of defence genes not initially captured in our analysis showed similar patterns for BIK1 and RBOHD during bacterial infection, in particular in the *atg5* background (Fig. 5a).

**Fig. 5:**
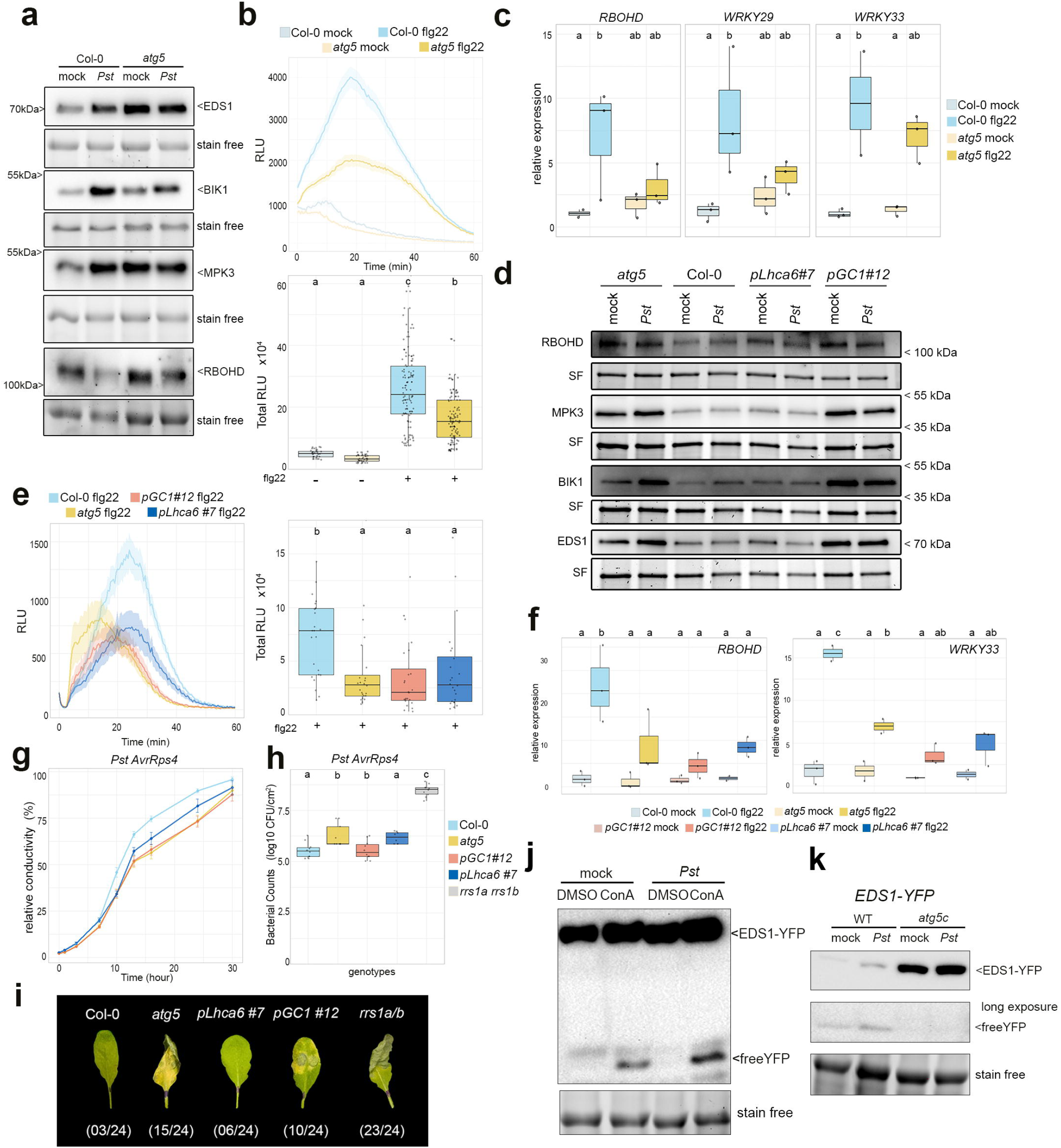
Autophagy plays a role in mesophyll-specific immune potentiation. **a**, Immunoblot analyses of immune components RBOHD, BIK1, EDS1 and MPK3 in Col-0 and *atg5* *A. thaliana* leaves 24 hours after syringe-infiltration with either 5×10^7^ CFUs/mL *Pst* or 10 mM MgCl2 mock. The large subunit of the Rubisco visible through Stain-Free imaging revelation serves as loading control. Representative results from at least three independent experiments. **b**, Reactive oxygen species (ROS) production measured by luminometry over 1 hour in leaves of 5-week-old Col-0 and *atg5* plants treated with 10 µM flg22 or water as mock. The upper panel shows ROS production kinetics expressed as relative light units (RLUs), with shaded areas indicating the standard error of the mean (SEM). The lower panel shows total ROS production over 1 hour measurement. Data from three independent experiments. **c**, Quantitative RT-PCR is shown for immune response marker genes (*RBOHD, WRKY29* and *WRKY33*) 60 min after 1 µm flg22 or water (control) treatment in 5-week-old Col-0 and *atg5 A. thaliana* plants. d, Immunoblot analyses of immune components RBOHD, BIK1, EDS1 and MPK3 in Col-0, *atg5* and *atg5*; *pLhca6::ATG5-GFP* and *atg5*; *pGC1:: ATG5-GFP A. thaliana* leaves 24 hours after syringe-infiltration with either 5×10^7^ CFUs/mL *Pst* or 10 mM MgCl2 mock. The large subunit of the Rubisco visible through Stain-Free imaging revelation serves as loading control. Representative results from at least three independent experiments. e, Reactive oxygen species (ROS) production measured by luminometry over 1 hour in leaves of 5-week-old Col-0, *atg5* and *atg5*; *pLhca6::ATG5-GFP* and *atg5*; *pGC1:: ATG5-GFP A. thaliana* plants. treated with 10 µM flg22. The left panel shows ROS production kinetics expressed as relative light units (RLUs), with shaded areas indicating the standard error of the mean (SEM). The right panel shows total ROS production over 1 hour measurement. Data from three independent experiments. f, Quantitative RT-PCR is shown for immune response marker genes (*RBOHD* and *WRKY33*) 60 min after 1 µm flg22 treatment or water (control) in 5-week-old in Col-0, *atg5* and *atg5*; *pLhca6::ATG5-GFP* and *atg5*; *pGC1::ATG5-GFP A. thaliana* plants. g, Electrolyte leakage over time in wild-type Col-0, *atg5, atg5*; *pLhca6::ATG5-GFP* and *atg5*; *pGC1:: ATG5-GFP* plants following syringe-inoculation with 2×10^8^ CFUs/mL *Pst* expressing AvrRps4. Conductivity is expressed as percentage of maximal conductivity after boiling. Lines represent means and error bars indicate standard error of the mean from 8 biological replicates of 6 leaf discs each. Similar results were obtained in two independent experiments. h, Bacterial population in 5-week-old *A. thaliana* leaves of Col-0, *atg5, atg5*; *pLhca6::ATG5-GFP, atg5*; *pGC1:: ATG5-GFP* and *rrs1a/rrs1b* plants syringe-inoculated with 2×10^5^ CFUs/mL *Pst* expressing AvrRps4. i, Representative images of disease symptoms observed in h. Numbers indicate symptomatic leaves out of the total number of infiltrated leaves. j, EDS1 degradation assay revealed by the ratio of free YFP to full-length EDS1-YFP. Immunoblot against GFP on crude extracts of 10-day-old *A. thaliana* seedlings EDS1-YFP in *eds1-2* background, infected with *Pst* or 10 mM MgCl_2_ mock for 8 hours following overnight 1 µm concanamycin A. k, EDS1 degradation assay revealed by the ratio of free YFP to full-length EDS1-YFP. Immunoblot against GFP on crude extracts of 10-day-old *A. thaliana* seedlings EDS1-YFP in *eds1-2* and *eds1-2 atg5* crispr background, infected with *Pst* or 10 mM MgCl_2_ mock for 8 hours. b,c,e,f,h Different letters indicate statistically significant differences, determined by one-way (b,e, h) or two-way ANOVA (c,f) followed by Tukey HSD post-hoc tests (adjusted p < 0.05).

We next hypothesized that immune responses are elevated in *atg5* and therefore re-assessed its capacity to mount PTI, given the enrichment of immune-associated genes. Unexpectedly, while MPK phosphorylation appeared to be enhanced in *atg5*, the flg22-induced ROS burst and expression of canonical PTI marker genes were compromised (Extended Data Fig.7a, Fig. 5b, c). In contrast, guard cells retained responsiveness to flg22, as stomatal closure remained largely intact (Extended Data Fig. 7b). A closer inspection of our scRNA-seq data revealed that, although guard cells exhibited a modestly elevated PTI score (based on^47^) in *atg5*, other cell types, including mesophyll and epidermal cells, did not show this increase (Extended data Fig. 7c,d). To investigate the contribution of cell-type-specific autophagy to immune responses, we leveraged our complementation lines. Mesophyll-specific autophagy restored immune protein levels to that of the wild-type but failed to complement ROS production or PTI gene expression (Fig. 5d-f), suggesting that PTI competence involves coordinated autophagy across tissues. By contrast, HR competence appears to be primarily mesophyll-autonomous as mesophyll-specific complementation significantly improved AvrRps4- and AvrRpm1-triggered HR compared with *atg5*, while the guard cell-specific complementation did not (Fig. 5g; Extended Data Fig. 7e). This is also reflected by the limitation of bacterial multiplication and disease progression of AvrRps4-delivering bacteria in Col-0 and mesophyll-complemented *atg5* (Fig. 5h,i).

Since AvrRps4- and, partially, AvrRpm1-triggered HR require EDS1^20^, we hypothesized that autophagic recycling of EDS1 is essential for robust immune activation. Using an *eds1*/EDS1-YFP reporter line^48^, we observed the release of free YFP upon ConA treatment being more prominent during bacterial infection (5j, Extended Data Fig. 7f). Free YFP was absent in CRISPR-inactivated *atg5c* plants^49^ infected with *Pst* (Fig. 5k) indicating that EDS1-YFP undergoes autophagic flux during bacterial infection. Consistent with this, selective autophagic degradation of EDS1 mediated by Nitrilase 1 (NIT1) has recently been described during viral infection^49^. Strikingly, our single-cell analysis reveals that *NIT1* expression is preferentially enriched in the disease-associated M8 cluster and reduced in *atg5*, whereas the related *NIT2* and *NIT4* are instead enriched in the immune-activated, *atg5*-specific M11 mesophyll population (Extended Data Fig. 7g). Together, these results suggest that autophagy spatially modulates EDS1 turnover across distinct mesophyll immune states, thereby contributing to state-specific differences in EDS1 accumulation and immune output.

## Discussion

Plant immunity requires the coordinated activation of defence pathways that are both spatially and temporally regulated to restrict pathogen growth while limiting collateral damage. Although major immune pathways are known to converge on shared signalling nodes^3,5,6^, how this coordination is achieved across different cell types remains unclear. Here, we demonstrate that autophagy acts as a central organizer of this coordination by partitioning immune strategies across distinct cell types during bacterial infection (Fig. 6). This spatial framework provides a unifying explanation for the previously reported, seemingly contradictory roles of autophagy in immunity-related cell death and susceptibility^20–23^.

**Fig. 6:**
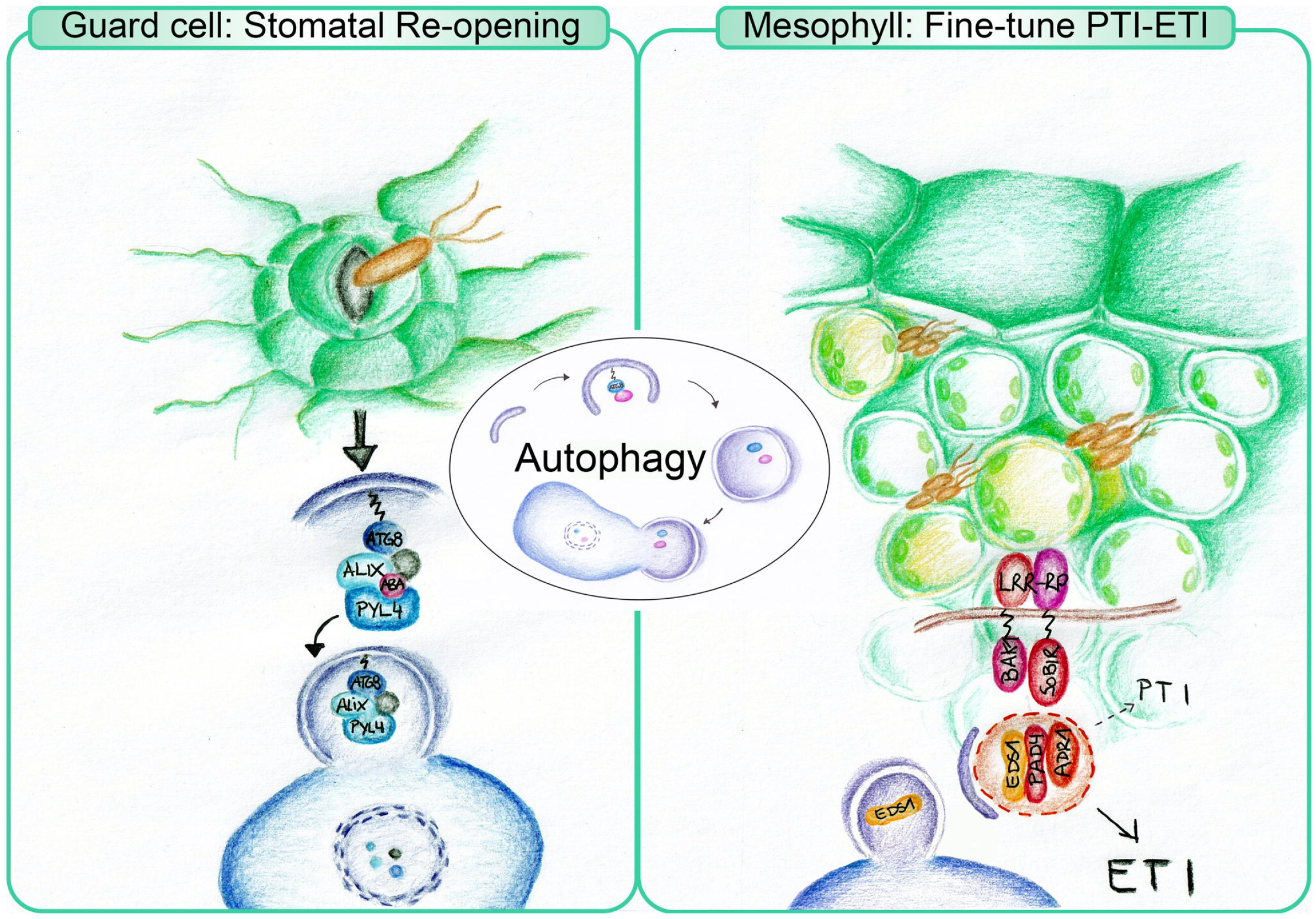
Autophagy spatially organizes plant immunity across cell types. Working model derived from scRNA-seq and functional analyses. Guard cells (left): Autophagy promotes stomatal reopening during infection by targeting the ABA receptor PYL4 for degradation via ATG8–ALIX, thereby suppressing ABA signalling. Mesophyll (right): Autophagy fine-tunes PRR signalling and likely enables effective ETI via the EDS1–PAD4–ADR1 node and EDS1 turnover.

Our data reveal that autophagy fulfils fundamentally different functions at the leaf surface and within internal tissues. In guard cells, autophagy promotes pathogen-induced stomatal reopening by facilitating the degradation of the ABA receptor PYL4, thereby enabling suppression of ABA signalling during infection. This finding extends previous work showing that stomatal immunity is tightly regulated by ABA signalling and actively manipulated by bacterial pathogens^16,31,50^ and provides a mechanistic explanation for why *Pseudomonas* activates autophagy during early stages of infection^22^.

In contrast, autophagy controls immune activation in mesophyll cells. Loss of autophagy leads to an enhanced expression of key defence regulators, including components of the EDS1-PAD4-ADR1 signalling module, in mesophyll cells, resulting in a primed immune state. As this module represents a central hub integrating immune signalling and promoting SA biosynthesis^5,51,52^, its misexpression suggests that autophagy constrains both the amplitude and spatial deployment of SA-dependent defence outputs. Consistent with recent single-cell studies revealing extensive heterogeneity in immune responses^19,44,53^, our data indicate that autophagy contributes to shaping these cell type–specific immune states. Together, these findings establish autophagy as a regulator of the spatial distribution and intensity of immune signalling rather than a simple positive or negative regulator of defence.

A key conceptual advance emerging from our study is the uncoupling of immune activation from execution. Despite elevated expression of central immune regulators, autophagy-deficient plants fail to mount robust EDS1-dependent hypersensitive cell death^20^. Our complementation analyses sharpen this picture: neither guard cell– nor mesophyll-specific restoration of autophagy rescue PTI responses, consistent with PTI competence being multicellular and potentially requiring coordinated autophagy across tissues. We hypothesize that this multicellular requirement reflects the central role of the EDS1-PAD4-ADR1 node in PTI-ETI potentiation^5,6^. Since productive PTI outputs may depend on the coordinated activity of this hub across multiple cell types, its spatial misregulation in autophagy-deficient tissue would selectively impair PTI competence. By contrast, mesophyll-specific complementation is sufficient to partially restore EDS1-dependent HR, establishing HR execution as a mesophyll-autonomous process. This dissociation suggests that the spatial requirements for PTI competence and HR execution are fundamentally distinct.

We propose that loss of autophagy disrupts the spatial organization and functional integrity of the EDS1-PAD4-ADR1 node, thereby limiting its capacity to trigger localized HR while leaving, or even amplifying, upstream signalling outputs intact. Sustained or misregulated EDS1 activity may continue to drive salicylic acid biosynthesis and systemic defence^50^, potentially explaining the paradoxical coexistence of enhanced immune signalling with impaired cell death and, under certain conditions, runaway responses^21^. Consistent with this, autophagy has been proposed to act as a molecular rheostat that constrains immune signalling by targeting EDS1 for degradation^49^. Strikingly, our study reveals the opposite during bacterial infection induced cell death: autophagic EDS1 turnover is required to promote, not restrict, HR execution, an apparent contradiction most likely explained by distinct cell-type-specific responses during bacterial infection and the different roles of distinct EDS1 complexes in SA and HR regulation. The cell-type-specific partitioning of nitrilase isoforms in our dataset raises the possibility that EDS1 turnover is differentially regulated across tissues, adding a spatial dimension to its regulation. Collectively, our findings reframe autophagy not merely as a brake on excessive immune activation, but as an active organizer of the spatial signalling architecture required for cell-type-appropriate immune execution.

Taken together, we propose a model in which autophagy coordinates plant immunity across cellular scales: promoting pathogen entry at the stomatal interface while simultaneously constraining and spatially organizing immune signalling in mesophyll cells. By linking proteostasis pathways to cell type–specific immune regulation, our findings provide a conceptual framework for understanding how plants balance defence activation with tissue integrity. Collectively, this work establishes autophagy as a critical spatial regulator of immune coordination and suggests that spatial control of signalling may represent a general principle underlying complex immune responses.

## Materials and Methods

### Plant material and growth conditions

*Arabidopsis thaliana* ecotype Col-0 wild-type and its derived T-DNA mutants and transgenic plants listed in Table S7. The double mutants *atg5 x ost1-3, atg5 x aba2-1*, atg5x *atg5 x pGC1*::ABAleon2.15 were obtained by crossing. The other transgenic lines were obtained by *Agrobacterium tumefaciens* floral dipping of Col-0 plants and the single insertion transgenics lines were selected and crossed with *atg5-1*.

Plants were grown under either short day or long day conditions (light/dark cycles: 16h 22°C/8h 20°C, 130 µmol·mm^2^·s^−1^ light intensity, 70% relative humidity) for *in vitro* experiments and seed propagation, or short day conditions (light/dark cycles: 12h 22°C/12h 20°C, 90 µmol·mm^2^·s^−1^ light intensity, 70% relative humidity) for bacterial infection assays. For *in vitro* experiments, seeds were sterilized with 70% ethanol and sown on solid 1% sucrose-supplemented ½ Murashige and Skoog (MS) medium or sucrose-free medium for carbon starvation assays. For infection assays, plants were grown for 4-5 weeks. Seeds for both experiments *in vitro* and with adult plants were stratified for 2 days at 4ºC. Nicotiana benthamiana wild-type were grown under long day conditions (light/dark cycles: 16h/8h, at 21°C and 70% humidity) for 4-6 weeks until use.

### Bacterial strains and growth conditions

*Pseudomonas syringae* pathovar *tomato* wild type strain DC3000 and its derived mutants listed in Table S8 were grown in King’s B (KB) medium supplemented with 100 mg/L rifampicin at 28ºC. *Agrobacterium tumefaciens* C58C1 derived strains listed in Table S8 were grown in Luria-Bertani broth (LB) medium supplemented with the appropriate antibiotics at 28ºC. *Escherichia coli* TOP10 and DB3.1 derived strains were grown in LB medium supplemented with the appropriate antibiotics 37°C.

### Molecular cloning

The list of plasmids used and generated in this study are listed in Table S9. The list of primers used to amplify the coding sequences from *A. thaliana* genes are listed in Table S10. For GreenGate based cloning, CDS or promoters were amplified with addition of flanking regions including BsaI sites and final constructs were generated using the CDS or promoters module of interest and desired other modules and assembled in the pGGP AG or pCAMBIA modified to be compatible with the GreenGate system^54,55^. For GATEWAY™ based cloning, CDS were amplified with addition of attb1/attb2 flanking regions and final constructs were generated through BP clonase™ II enzymatic reaction and LR Clonase™ II enzymatic reaction into the desired destination vectors. All entry clones were verified by sanger sequencing.

### Plant inoculation and bacterial growth assays

Overnight *Pst* cultures were centrifuged for 7 minutes at 4,000 rpm. Pellets were washed twice with 10 mM MgCl_2_ prior to measuring the optical density at 600 nm (OD_600_). For bacterial growth assays after dipping, bulk or scRNA transcriptomic, and stomatal aperture, *Pst* inocula were prepared at a final OD_600_ of 0.2 in 10 mM MgCl_2_ and 0.02% Silwet L-77. For bacterial growth assays after mesophyll infiltration with wild type or *avrRps4*-containing *Pst* strains, inocula were prepared at a final OD_600_ of 0.0001 and 0.0005 respectively. Plants were fully dipped in the bacterial solution for 1 minute. For bacterial growth assays upon syringe inoculation, the final OD_600_ was adjusted to 0.0001 in 10 mM MgCl_2_.The bacterial inocula were infiltrated with a needleless syringe into the abaxial side of fully developed leaves from 5-6-week-old *A. thaliana* plants. For immune reporter lines, *Pst* Δ*hrcC* inocula were prepared at a final OD_600_ of 0.1 in 10 mM MgCl_2_ and infiltrated with syringe. After inoculation, all plants were well-watered and put in covered trays to reach nearly saturated humidity conditions. Unless stated otherwise, plants were used for experiments 2 days post inoculation for bulk or scRNA transcriptomic, 3 days post inoculation for bacterial growth assays, 3 hours after inoculation for stomata aperture, 24hours after inoculation for immune reporter lines. Bacterial populations were estimated by plating serial dilutions of bacteria extracted from two 6-mm-diameter leaf discs in 200 µL 10 mM MgCl_2_ homogenized with a tissue lyser (Retsch). Colony forming units were manually counted 48 hours after plating. For liquid inoculation, 10-day-old *A. thaliana* seedlings were incubated with bacterial solution at a final OD_600_ of 0.2 in 10 mM MgCl_2_ for 2 or 5 hours until use.

### Laser scanning confocal microscopy and image analysis

Images were taken with a confocal laser scanning microscope (STELLARIS 8; Leica) using a water-immersed objective (HC PL APO CS2 20× or 40x /1.20 W) with a resolution acquisition set to 512 x 512 (reporter line analyses) or 1024 × 1024 (for subcellular localization analyses) and a line average of 4. The following excitation and emission settings were used: GFP (488 nm excitation, 500–560 nm emission), YFP (514 nm, 530–570 nm), RFP/mCherry (587 nm, 600– 640 nm), and mTurquoise (448 nm, 460–550 nm). Tau-gating was used to remove autofluorescent signal. Chlorophyll autofluorescence was visualized in the far-red wavelength. For leaves expressing the pGC1::ABAleon2.15 construct were fixed with formaldehyde 4% prior imaging. For all the reporter lines, Z-stack images were acquired with a fixed step size adjusted per experiment to cover the entire range of signal along the z-axis. Image processing was performed using ImageJ;. For clarity purposes, contrast on the individual channels was manually adjusted.

### Quantification of confocal images

Nuclear GFP, YFP or mTurquoise signals were quantified using a custom ImageJ macro. For each image series, the GFP, YFP or mTurquoise channel was extracted, and a maximum intensity Z-projection was generated. An automatic threshold (Li or MaxEntropy method for mesophyll and IsoData method for epidermis and guard cells) was applied to create a binary mask, followed by morphological opening (circle, radius 2.5) to refine nuclei outlines. Particles were analyzed to generate regions of interest (ROIs). For genes expressed in nuclear of both epidermis and guard cells, each ROI was manually verified via a dialog prompt to confirm it represented nucleus from a guard cell or pavement cell. For *pATG8c::GFP-GUS* and *pGC1*::ABAleon2.15, ROIs were manually drawn over guard cells. For confirmed nuclei or cells, mean intensity was recorded. Between 100 to 500 cells from 4-8 leaves were measured for each data point.

### RNA sequencing

Sequencing libraries were prepared from the extracted RNA using the NEBNext® UltraTM II RNA Library Prep Kit for Illumina® and Dual Index Primers (New England Biolabs) following the manufacturer instructions. The quality of the libraries was verified with a Bioanalyzer (Agilent) prior to send for Illumina® sequencing at the NovaSeq PE150 Platform of Novagene (Cambridge, UK). After quality control, transcript abundances were quantified using Salmon and aggregated to gene-level counts with tximport based on the *A. thaliana* TAIR10.54 reference genome.The log_2_ of the fold change (log2FC) and false discovery rate (FDR) were determined using R package DEseq2 with default settings^56^. A gene is considered significantly differentially expressed when the absolute value of its log2FC > 2 and its FDR < 0.05.

### Protoplast isolation and ScRNA sequencing

For protoplast isolation, tape-sandwich method was modified and applied^57^, and a subset of leaves was processed by cutting and finely dicing the tissue with a scalpel blade. For both methods, the abaxial epidermis, guard cells, and trichomes and the adaxial side of the fully developed leaves were incubated in digestion buffer (2% Cellulase-RS, 0.4% Macerozyme R10, 2 mM MES pH 5.7, 0.6 M mannitol, 10 mM KCl, 10 mM MgCl_2_, 0.1% BSA) for 1 h at room temperature with gentle agitation^58^. The mixture was subsequently filtered through a 70 μm nylon filter and rinsed with protoplast buffer (2 mM MES pH 5.7, 0.6 M mannitol, 10 mM KCl, 10 mM MgCl_2_, 0.1% BSA). After low-speed centrifugation (200g, 2 min, room temperature), the pellet was resuspended in protoplast buffer. This wash procedure was repeated in total 3 times, the protoplasts centrifuged as before and resuspended in ~700 μl or less protoplast buffer and filtered through a 40 μm cell strainer (Flowmi Bel Art SP Scienceware). Protoplasts were validated under a light microscope and quantified using a haemocytometer and adjusted to a density of approximately 800-900 cells per μl. For each sample, twelve to thirteen leaves were used.

Single cell RNA-seq libraries were prepared from fresh protoplasts according to the 10x Genomics Next GEM Single Cell 3’ Reagent Kits v3.1 protocol. ScRNA-seq library sequencing was performed on the NextSeq (Illumina) platform.

### ScRNA-seq analysis

### Raw data processing and quality control

Sequence data were processed to obtain single-cell feature counts by running Cell Ranger (v.6.1.2) for scRNA-seq data and the reads were aligned to *Arabidopsis* TAIR10 reference genome, which was downloaded from https://plants.ensembl.org/. Count data were analysed using the R packages Seurat (5.2.1). Quality control filtering was performed to retain cells with mitochondrial gene content below 1%, chloroplast gene content below 10%, number of detected genes between 300 and 6,000, and total UMI counts between 500 and 40,000.

#### Integration and clustering

To mitigate the effects of protoplasting, previously identified protoplasting-induced genes^57^ were removed from the dataset for clustering analysis. Cell cycle variation was regressed out using cell cycle genes^59^ (Zhang et al. 2021). Before integration, doublets were removed using DoubletFinder^60^.

Data were normalized with SCTransform (v2) and integrated across samples via RPCA-based anchor identification (k.anchor = 5). After integration, the data were normalized again using NormalizeData on both the protoplasting-filtered and the full RNA assays to facilitate downstream gene expression analysis. Clustering was performed on the integrated object with PCA (1:50), graph-based clustering (resolution = 0.5), and UMAP visualization (reduction = “pca”, dims = 1:50).

To integrate our dataset with published dataset (GSE213622)^17^, we also use RPCA-based anchor integration after SCTransform normalization. Clustering was performed on the integrated object (PCA 1:50, resolution = 0.7).

#### Identification of markers, DEGs and heat map generation

Cluster-specific marker genes that are conserved across samples were identified using the FindAllMarkers function in Seurat, with parameters grouping.var = “orig.ident”, only.pos = TRUE and logfc.threshold = 0.25. Genes with an adjusted p-value < 0.01 were retained as markers, corresponding to genes enriched in the target sample relative to all others.

Cell-type-specific DEGs between *atg5*-mock and Col-0-mock samples was identified using the FindMarkers function (log_2_FC ≥ 0.5, min.pct = 0.25) in Seurat. Mock samples from 2 dpi and 3 hpi were pooled into *atg5*-mock and Col-0-mock groups. Genes with log_2_ fold change ≥ 0.5 and detected in at least 25% of cells in either group were considered for analysis.

DEGs enriched in M11 versus other mesophyll cells at 2 dpi were identified using FindMarkers (log_2_FC ≥ 0.5, min.pct = 0.25) separately in Col-0-Pst-2dpi (genes upregulated by *Pst*) and atg5_mock_2dpi (genes upregulated by *atg5* mutation) samples. Genes with an adjusted p-value < 0.05 were considered significantly differentially expressed. Overlapping upregulated genes were considered as atg5-specific immune genes.

Genes upregulated in M8 versus other mesophyll cells in Col-0-Pst-2dpi were overlapped with genes downregulated in M8 of atg5_mock_2dpi versus Col-0-Pst-2dpi. The overlapping set was defined as atg5-specific disease genes (log_2_FC ≥ 0.5, min.pct = 0.25, p_val_adj < 0.05).

DEGs between guard cell subtypes GC20 and GC16 were identified using FindMarkers (log_2_FC ≥ 0.5, min.pct = 0.25, min.diff.pct = 0.25, p_val_adj < 0.05). Genes with avg_log_2_FC > 2 were defined as highly expressed in GC20.

For the heatmaps of immune-, disease-related genes (Fig 1F), cell-type-specific atg5-upregulated genes (Fig S1h), ATGs (Fig 1A), atg5-specific immune genes (Fig 4B), PTI genes (Fig 4G), upregulated by *Pst* genes(Fig S5i), atg5-specific disease genes (Fig S6C), the Seurat object was subset to the cell types and conditions shown in each figure. Pseudobulk expression was obtained with AggregateExpression (assay = “RNA”) and Heatmaps were generated with row-scaled scale.data.

#### Signature score computation

A Pst_responsive score was computed previously described^17^, using AddModuleScore (Seurat), defined as the score for *Pst*-upregulated genes minus the score for *Pst*-downregulated genes (from bulk RNA-seq data).PTI score was calculated via AddModuleScore using PTI gene sets from^47^.

##### GO enrichment

Gene Ontology enrichment was performed using the topGO R package. Gene-to-GO annotations for biological process (BP) were retrieved from Ensembl Plants (BioMart). The background gene set comprised all *Arabidopsis* genes from the Ensembl annotation. For a given set of genes of interest, enrichment was assessed using Fisher’s exact test with the weight01 algorithm to account for GO hierarchy. GO terms with a corrected p-value < 0.05 were considered significantly enriched. Results were visualised as a dot plot showing the top terms, with point size representing the number of significant genes and colour indicating −log_10_(p-value).

### STOMATAL aperture measurement and analysis

5-6-week-old *A. thaliana* plants were kept for at least 3 hours after lights on to ensure stomata open before starting experiments. The plants were dip-inoculated with Pst or the detached leaves were immersed in 10 mM MgCl_2_ with Pst or mock(10 mM MgCl_2_). For ABA and Fluridone treatment, 30 μM ABA or 50 μM Fluridone or mock (0.1% ethanol) spraying was performed 1 hour after Pst infection. For flg22 treatment, epidermis was peeled and immersed in buffer with 5 μM flg22 or mock (water). The leaves were harvest at desired time and immediately immersed in stomatal fixation solution (formaldehyde 4%) for 1 min to stop stomatal movement^16^. The leaves were then cleaned by overnight incubation in 9:1 ethanol:acetic acid solution, followed by a series of ethanol washes (from 70%, 50%, down to 20%) leaves were stained with 5 μM rhodamine 6G for 1 min and were imaged using a Leica M205C stereomicroscope. Stomatal aperture was quantified using a custom ImageJ macro. For each image, two perpendicular lines were manually drawn per stoma using the line tool: one along the stomatal length (major axis) and one across the pore (minor axis). The ROI Manager recorded the selections, and a multi-measure function extracted the length of each line. The aperture ratio was calculated as minor axis /major axis. from 100 to 400 stomata from 4-8 leaves were measured for each data point.

### Carbon starvation and pigment measurements

10-day-old *A. thaliana* seedlings were wrapped with Aluminum foil and kept in darkness for 11 days. For photosynthetic pigment concentration, 3 rosettes were sampled and incubated over-night in 1mL acetone 100% on a rotating wheel in cold room. Tubes were next centrifuged for 3min at 400g. 200µl of was then pipetted in a transparent 96-well plate and absorbance was measured at 470nm, 646nm and 663nm using a plate reader (Tecan Infinite 200 PRO®). Acetone 100% was used as blank. Concentration was calculated as following the obtained from Karlsruher Institut für Technologie (KIT, https://www.jkip.kit.edu/molbio/998.php, “Chlorophyll and Carotenoid determination in leaves”). Values were then then expressed as ng.mm^2^.

### Immunoblotting

Proteins were extracted in extraction buffer (100 mM Tris pH 7.5, 1 mM EDTA, 3% SDS) and mixed with 4X Laemmli buffer (BioRad) prior to boiling at 95ºC for 10 minutes and clearing by centrifugation of 1 minute at 13,000 rpm. In case of PYL4, protein was extracted with buffer (50 mM Tris-MES pH 7.4, 0.5 M Sucrose, 1 mM MgCl_2_, 10 mM EDTA, 5 mM dithiotreitol, 0.2% (v/v) Nonidet P-40). Total protein extracts were separated by SDS–PAGE, visualized with the Stain-Free Imaging Technology in a ChemiDoc Imaging System (BioRad), transferred to PVDF membranes (BioRad), blocked with 5% skimmed milk in TBS (200 mM Tris and 1500 mM NaCl), for 1 hour at room temperature and incubated with the correspondent antibody listed in Table S11 for 1-2 hours at room temperature or overnight at 4ºC. For the detection, the immunoreaction was developed using an ECL Prime Kit (GE Healthcare) and detected with a ChemiDoc Imaging System (BioRad). When the Stain-Free images of the total protein extracts were not taken, Ponceau S staining (0.1% Ponceau Red and 5% acetic acid) was performed on the membranes after revelation instead. The relative quantification of the protein band intensities and ratios was performed using the Image Lab software (BioRad).

### Transient expression in *Nicotiana benthamiana* leaves

Overnight *A. tumefaciens* cultures were centrifuged for 7 min at 4,000 rpm. Pellets were washed twice with agroinfiltration buffer (10mM MgCl_2_, 10mM MES pH 5.7) prior to measuring OD_600_. Inocula were prepared at a final OD_600_ of 0.1-0.5 in 200µM acetosyringone-supplemented agroinfiltration buffer and incubated at room temperature and darkness for 2 hours before infiltration with a needleless syringe into the abaxial side of 4 to 6-week-old *N. benthamiana* leaves. Plants were used for further experiments 24-48 hours after infiltration.

### Co-immunoprecipitation

Plant tissue was ground with a mortar and liquid nitrogen and then extracted in 1ml/g of GTEN buffer (10% glycerol, 25 mM Tris pH 7.5, 1 mM EDTA, 150 mM NaCl, 1 mM DTT, 1X Protease inhibitor Cocktail (Merck) and 0.5% Triton X,). The solution was vortexed until homogenized and incubated at 4ºC with rotation for 10min following centrifugation for 30 minutes at 4000 g and 4°C. Supernatant was filtered with three layers of Miracloth (Merck). 50µl of the filtrate was sampled as input, supplemented with 12.5µL Laemmli buffer 4X (BioRad) and boiled for 10 minutes at 95°C. Next, 10µl/g of GFP-Trap® Agarose beads (ChromoTek) were added to the remaining filtrate and incubated for 2 hours at 4ºC. The filtrates were then centrifuged at 800 g for 1 minute, the supernatants were carefully removed, and the pelleted beads were transferred to new 1.5mL microcentrifuge tubes in 1mL of GTEN buffer. Five steps of washing were performed by centrifuging at 800 g for 1 minute using 1mL GTEN buffer. After the last washing, 2X Laemmli buffer (BioRad) was added to equal amounts of beads and the samples were boiled 10min at 95°C prior to immunoblotting as previously described.

### Measurement of ROS production

16 to 24 4-mm-diameter leaf discs were harvested for each condition and individually incubated overnight in a 96-well plate (PierceTM; Thermo Scientific) containing 100 μL of distilled water. For ROS production measurements, water was replaced by 100 μL of elicitor mix (10 μM flg22, 100 μM luminol, 20 μg/mL horseradish peroxidase) or mock (elicitor mix without flg22) and relative luminescence was quantified in each well every minute during 1 h, using a plate reader (Tecan). The total amount of luminescence per condition was calculated as the average of the sums of all measurements taken during the recorded hour per sample.

### RNA extraction and quantitative real-time PCR

Total RNA from plant samples was extracted following the manufacturer instructions with either the RNeasy Plant Mini Kit (Qiagen) for RNA sequencing, or with the NucleoZOL reagent (Macherey-Nagel) for the rest of applications. After extraction, 500 ng of total RNA per samples were treated with DNase I (Thermo Scientific) prior to the cDNA synthesis using the LunaScript® RT SuperMix Kit (New England Biolabs), in both cases following the manufacturer instructions. Quantitative real-time PCR was performed using 2X MESA Blue qPCR Master Mix (Eurogentec) using a 2-step protocol for 40 cycles and melting curve analysis. The relative gene expression was then calculated following the ΔΔCt method. For the ribosomal enrichment, the relative expression for each condition/fraction was calculated separately, then the enrichment was calculated normalizing each ribosomal fraction to its respective total fraction. The primers used for qPCR are listed in Table S10.

### Electrolyte leakage assay

Electrolyte leakage assays were performed using 4-week-old *Arabidopsis thaliana* plants infiltrated in the mesophyll with *Pseudomonas syringae* pv. *tomato* strains expressing *avrRps4* or *avrRpm1* at final OD_600_ values of 0.5 and 0.05, respectively. Immediately after inoculation, 36–48 leaf discs (6 mm diameter) were harvested from infiltrated leaves and washed in double-distilled water for 15 min prior to distribution into 6–8 biological replicates consisting of six leaf discs each in 4 mL double-distilled water in 14 mL polypropylene graduated culture tubes (Simport). Conductivity was measured at the indicated time points using a FiveGo conductivity meter (Roth). Maximum conductivity was determined after the final time point by boiling the samples at 95°C for 10 min.

## Acknowledgements

We thank Margot Smit, Rainer Waadt, Vicente Rubio, Guillaume Née, Julien Gronnier, Jane Parker and Marion Clavel for providing published material. We are thankful to Lea Röhder for her technical support. We are thankful to Thorben Krüger for insightful discussions on ABA and ABA-related pathways. We are thankful to Paul Gouguet for stimulating discussions. We thank the master and bachelor students from the past three years of our course “Plant Cell Biology meets Plant–Pathogen Interactions,” who generated preliminary data inspiring this project. This work was supported by European Union’s Horizon 2020 research and innovation programme grant number 948996 DIVERSIPHAGY (S.Z. and S.Ü.), and funds from DFG GZ: INST 213/1180-1 FUGG (microscopy facility RUB).

## Author contribution

S.Ü. conceived and designed the project. S.Z. performed most of the experiments, analysed data and performed all computational studies. M.G.F, O.L., N.A., K.X. and N.S. performed experiments. G.L. helped in computational analysis. T.D., P.S. and M.T helped to setup the scRNA-seq pipeline. Y.D. and Q.F provided unpublished resources. S.Ü. wrote the manuscript together with S.Z..

