## Extended Data Figure 1-7 for "Autophagy acts as a spatial organizer of cell-type-specific plant immunity"

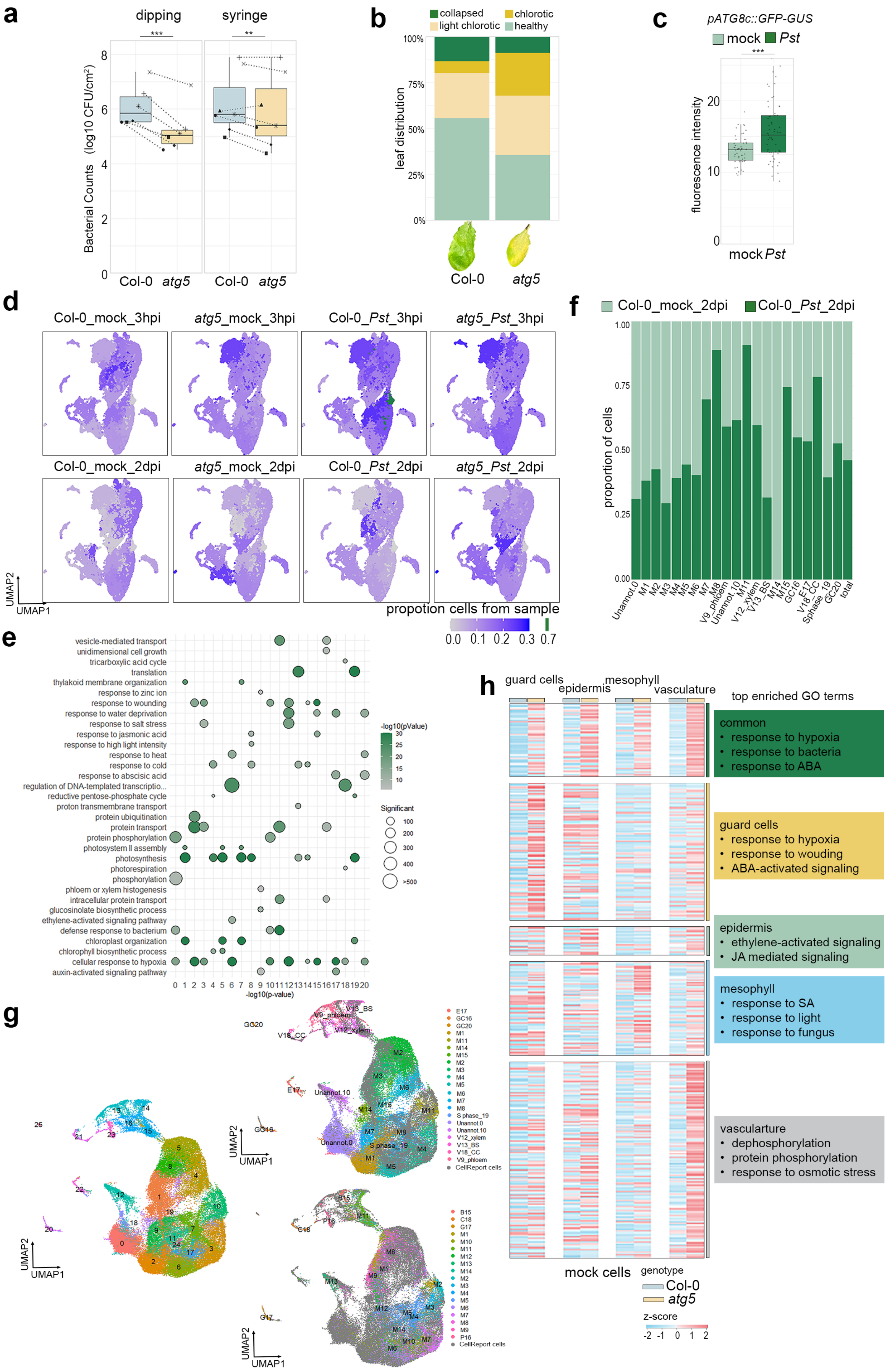

a
dipping
syringe
***
**
Bacterial Counts (log10 CFU/cm²)
Col-0 atg5
Col-0 atg5
b
collapsed
chlorotic
light chlorotic
healthy
leaf distribution
100%
75%
50%
25%
0%
Col-0
atg5
c
pATG8c::GFP-GUS
mock
Pst
***
fluorescence intensity
mock Pst
d
Col-0_mock_3hpi
atg5_mock_3hpi
Col-0_Pst_3hpi
atg5_Pst_3hpi
Col-0_mock_2dpi
atg5_mock_2dpi
Col-0_Pst_2dpi
atg5_Pst_2dpi
UMAP2
UMAP1
propotion cells from sample
0.0 0.1 0.2 0.3 0.7
f
Col-0_mock_2dpi
Col-0_Pst_2dpi
proportion of cells
Unannot.0 M1 M2 M3 M4 M5 M6 M7 M8 V9_phloem Unannot.10 M11 V12_xylem V13_BS M14 M15 GC16 E17 V18_CC Sphase_19 GC20 total
e
vesicle-mediated transport
unidimensional cell growth
tricarboxylic acid cycle
translation
thylakoid membrane organization
response to zinc ion
response to wounding
response to water deprivation
response to salt stress
response to jasmonic acid
response to high light intensity
response to heat
response to cold
response to abscisic acid
regulation of DNA-templated transcriptio...
reductive pentose-phosphate cycle
proton transmembrane transport
protein ubiquitination
protein transport
protein phosphorylation
photosystem II assembly
photosynthesis
photorespiration
phosphorylation
phloem or xylem histogenesis
intracellular protein transport
glucosinolate biosynthetic process
ethylene-activated signaling pathway
defense response to bacterium
chloroplast organization
chlorophyll biosynthetic process
cellular response to hypoxia
auxin-activated signaling pathway
-log10(pValue)
Significant
100
200
300
400
>500
-log10(p-value)
g
UMAP2
UMAP1
h
guard cells
epidermis
mesophyll
vasculature
top enriched GO terms
common
• response to hypoxia
• response to bacteria
• response to ABA
guard cells
• response to hypoxia
• response to wouding
• ABA-activated signaling
epidermis
• ethylene-activated signaling
• JA mediated signaling
mesophyll
• response to SA
• response to light
• response to fungus
vasculature
• dephosphorylation
• protein phosphorylation
• response to osmotic stress
mock cells
genotype
Col-0
atg5
z-score
-2 -1 0 1 2

**Extended Data Fig. 1: Autophagy acts in a cell-type specific manner**

**a**, Bacterial population in *A. thaliana* leaves of wild type Col-0 and *atg5* plants dip-inolucated with $10^8$ CFUs/mL *Pst* or syringe-infiltrated with 5 x$10^4$ CFUs/mL *Pst*. Data represent seven independent experiments, with each shape corresponding to an individual experiment. Significant differences between genotypes within each inoculation method are determined using two-way ANOVA with experiment as a random effect, followed by a Tukey HSD post-hoc test (adjust $^{**}p < 0.01$, $^{***}p < 0.001$). A significant interaction between genotype and inoculation method interaction was also detected ($p = 0.008$).

**b**, Leaf phenotypic analysis of the syringe-infiltration plants from (a). Stacked bar chart illustrates the distribution of leaf states (collapsed, chlorotic, light chlorotic, healthy) as percentages of total leaves per genotype. Data from six independent experiments (n = 90 leaves per genotype). The p value was calculated using a chi-square test ($\chi^2 = 30.72$, df = 3, $p = 9.7 \times 10^{-7}$).

**c**, GFP fluorescence intensity from guard cells of *A. thaliana* leaves expressing *pATG8c::GFP-GUS* at 24 hours upon flooding with $10^8$ CFUs/mL *Pst* or 10 mM $MgCl_2$ mock. Statistical differences between genotypes are assessed with Student's t-test ($^{***}p < 0.001$).

**d**, UMAP visualization of single cells colored by sample-specific cell type proportions. Each panel shows the same dimensional reduction embedding of integrated single-cell transcriptomes, with cells colored according to the proportion of cells from the given sample within their assigned cell type. Color intensity reflects the relative contribution of each sample to a cell type cluster, enabling visual comparison of cell type representation across genotypes, treatments, and time points.

**e**, GO enrichment analysis for marker genes of each cluster.

f, Stacked bar plot displaying the proportion of cells originating from the two samples (Col-0 mock 2d and Col-0 Pst 2d) within each cell type cluster.

**g**, UMAP visualization of integration of our dataset and the published *Pst* infection dataset[49]. The three panels display the same UMAP embedding colored by different annotations. Left: Integrated clusters identified in the combined analysis; Top right: Cell type annotations from this study; Below right: Cell type annotations from the published dataset.

**h**, Cell-type specific expression patterns of *atg5*-upregulated genes under mock conditions. Heatmap showing the scaled relative expression of the genes upregulated in *atg5* compared to Col-0 (mock). Genes are grouped into those commonly upregulated in all four cell types (guard cells, epidermis, mesophyll, vasculature) or specifically in a single cell type. Top GO terms for each group are listed on the right.

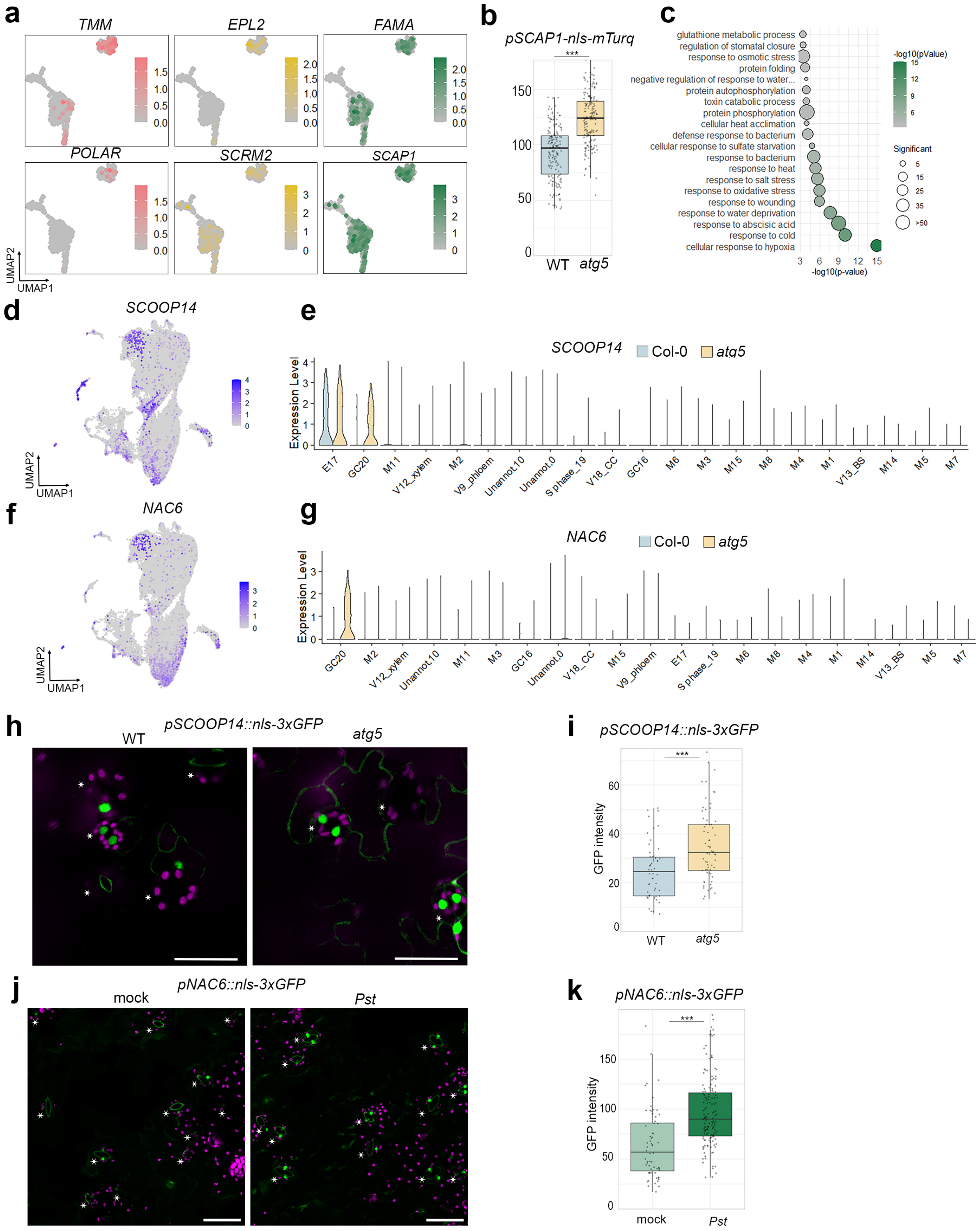


**Extended Data Fig. 2: Autophagy deficiency results in a specific guard cell cluster**

**a**, Expression patterns of guard cell development markers in two guard cell clusters.

**b**, mTurquoise fluorescence intensity from guard cells in cotyledons of 10-day-old *A. thaliana* expressing *pSCAP1::nls-mTurquoise2* in wildtype and *atg5* background.

**c**, GO enrichment analysis for the genes highly expressed in GC20 compared to GC16

**d,f** UMAP showing expression pattern of *SCOOP14* (d) and *NAC6* (f) in ScRNA data

**e,g** Violin plots showing expression of *SCOOP14* (e) and *NAC6*(g) across cell types in Col0 and *atg5*

**h**, Representative confocal images of 10-day-old *A. thaliana* leaves of *pSCOOP14::nls-3xGFP* in wild type and *atg5*

**j**, Representative confocal images of 10-day-old *A. thaliana* leaves of *pNAC6::nls-3xGFP* incubated with $10^8$ CFUs/mL *Pst* or mock for 6 hours. Fluorescent images are maximal intensity projections of Z-stacks.

**i,k**, quantification form (h) and (g), respectively. . This experiment was repeated three times with similar results.

**h,j**, star indicate the stomata positions. Scale bars = 50 µm. b,j,k, Statistical differences between genotypes are assessed with Student's t-test (****p* < 0.001).

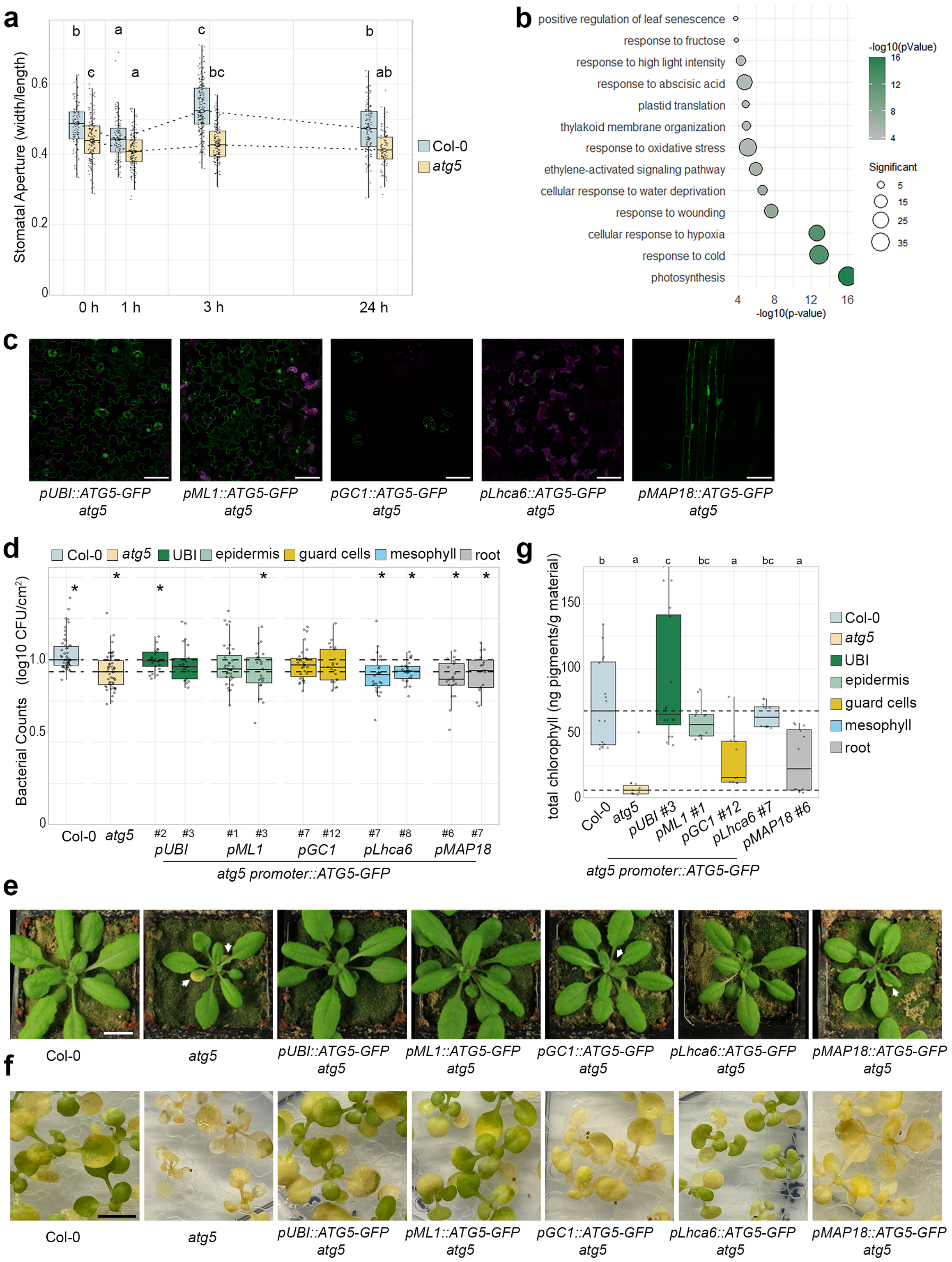

**Extended Data Fig. 3: Autophagy dependent guard-cell specific response during infection**

**a**, Stomatal apertures of 5-week-old *A. thaliana* Col-0 and *atg5* plants were measured at the indicated time points after dip-inoculation with $10^8$ CFUs/mL *Pst*. Different letters indicate statistically significant differences between time points within each genotype, determined by two-way ANOVA followed by Tukey HSD post-hoc tests (adjusted $p < 0.05$). Data from two independent experiments.

**b**, GO enrichment analysis for the M14 marker genes.

**c,** Confocal images of *atg5* complementation lines carrying tissue-specific promoters. Scale bar = 50 µm

**d**, Relative bacterial population in leaves of Col-0, *atg5* and *atg5* complementation lines carrying tissue-specific promoters. 5-week-old *A. thaliana* plants were dip-inoculated with $10^8$ CFUs/mL *Pst.* Log CFU values were normalized to the median of Col-0 for each independent experiment. Dashed horizontal lines indicate the overall medians of Col-0 and *atg5* across all experiments. one-way ANOVA test with experiment as random effect revealed a significant overall effect of genotype ($p < 0.001$). Post-hoc pairwise comparisons against Col-0 and against *atg5* were performed using the "mvt" correction for multiple testing. Significant differences (adjusted $p < 0.05$) are indicated by stars (*). Data from seven independent experiments.

**e**, Representative images of 5-week-old *A. thaliana* plants Col-0, *atg5*, and *atg5* complementation lines carrying tissue-specific promoters, grown under long-day conditions. Arrows indicated leaves showing early senescence. Scare bar represents 1 cm.

**f**, Representative images of *A. thaliana* seedlings Col-0, *atg5*, and *atg5* complementation lines carrying tissue-specific promoters, after 11days carbon starvation treatment. Scare bar represents 0.5 cm.

**g**, Total chlorophylls measurement of Fig. S3e. Dashed horizontal lines indicate the medians of Col-0 and *atg5* across all experiments. Different letters indicate statistical groups determined using one-way ANOVA with experiment as a random effect, followed by a Tukey HSD post-hoc test (adjusted $p <$ 0.05). Data from three independent experiments.

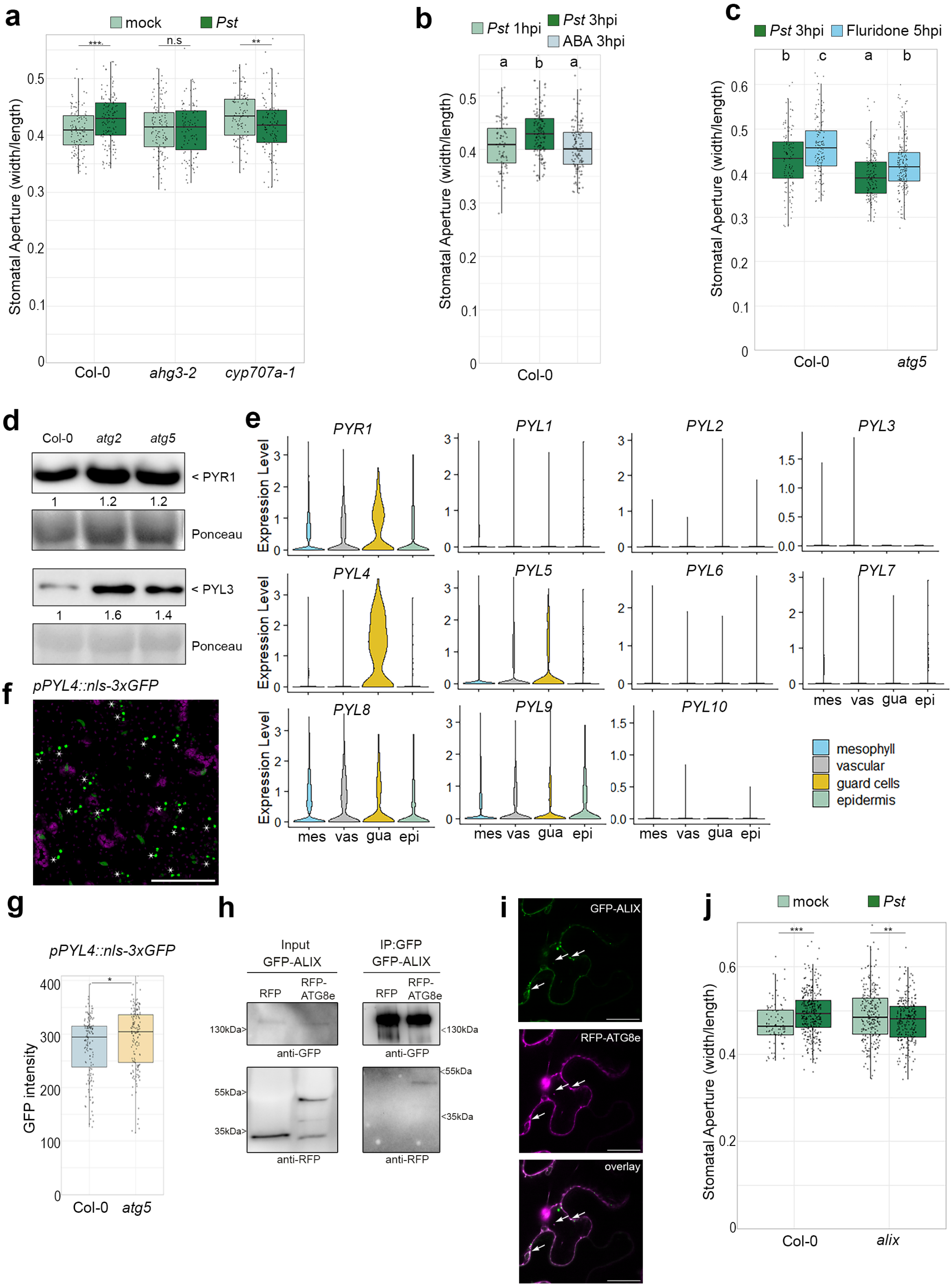


**Extended data Fig. 4 Autophagy influences stomatal opening by altering ABA activity through recycling of ABA receptor**

**a**, Stomatal apertures of 5-week-old *A. thaliana* Col-0, *ahg3-2* and *cyp707a-1* plants at 3 hours after dip-inoculation with $10^8$ CFUs/mL *Pst* or 10 mM $MgCl_2$ mock.

**b**, Stomatal response to ABA in *Pst*-infected Col-0 plants. 5-week-old *A. thaliana* plants were dip-inoculated with $10^8$ CFUs/mL *Pst* and then spayed with 30 µM ABA or ethanol as mock at 1hpi. Leaves were harvest and fixed at indicated time point.

**c**, Stomatal response to Fluridon in *Pst*-infected Col-0 and *atg5* plants. 5-week-old *A. thaliana* plants were dip-inoculated with $10^8$ CFUs/mL *Pst* and then spayed with 50 μM Fluridon or ethanol as mock at 1hpi. Leaves were harvest and fixed at indicated time point.
**b,c** Different letters indicate statistically significant differences, determined by one- (b) or two-way(c) ANOVA followed by Tukey HSD post-hoc tests (adjusted $p < 0.05$).
**d**, Immunoblot analysis of PYR1 and PYL3 protein levels in 10-day-old *A. thaliana* seedlings of Col-0, *atg2* and *atg5*. Stain-Free imaging revelation or Ponceau staining serve as loading control. Numbers indicate the band intensity of PYR1 or PYL3 immunoblot normalized to each sample loading. The experiment was repeated two times with similar results.
**e**, violin plots showing the expression pattern of detected *PYLs* genes across cell types in ScRNA data.
**f**, Confocal microscopy pictures of *A. thaliana* leaves expressing *pPYL4::nls-3xGFP*. stars point the stomata positions. Scale bars = 100 μm
**g**, Quantification fluorescence intensity from guard cells in cotyledons of 10-day-old *A. thaliana* expressing *pPYL4::nls-3xGFP* in wildtype and *atg5* background. Significant differences between genotypes are accessed using Student's t-test ($*p < 0.05$).
**h**, Coimmunoprecipitation of RFP or RFP-ATG8e with GFP-ALIX transiently expressed in *Nicotiana benthamiana.*
**i**, Confocal microscopy pictures of transiently co-expressed RFP-ATG8e with GFP-ALIX in N. benthamiana. White arrows indicate co-localization of RFP and GFP signals. Scale bars represent 25 μm.
**j**, Stomatal apertures of 5-week-old *A. thaliana* Col-0, and *alix* plants at 3 hours after dip-incubated with $10^8$ CFUs/mL *Pst* or 10 mM $MgCl_2$ mock.
a.j, Significant differences between treatment within each genotype are determined by two-way ANOVA followed by Tukey HSD post-hoc tests (adjusted $**p < 0.01$, $***p < 0.001$, n.s, not significant).

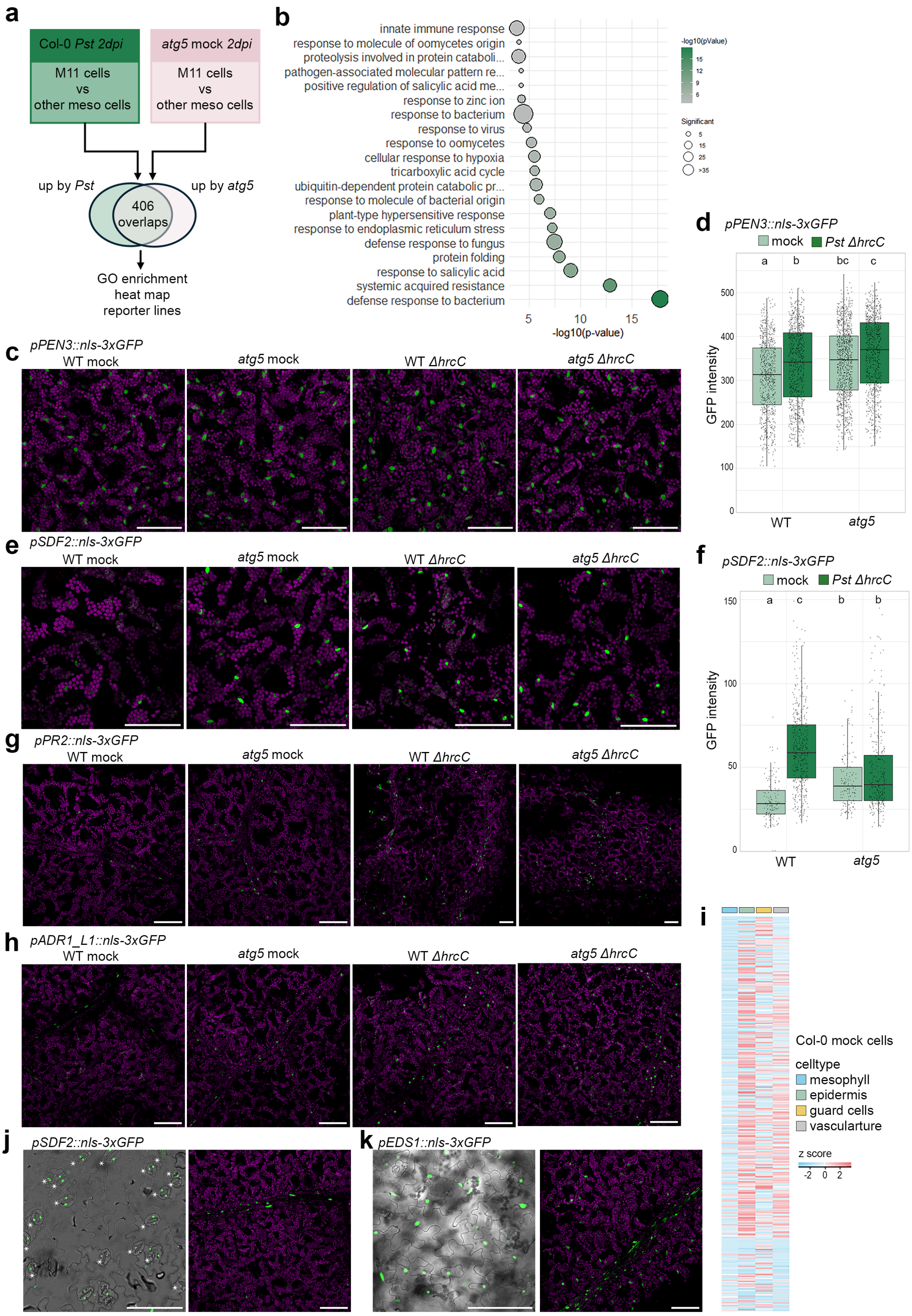


**Extended Data Fig. 5: Autophagy deficiency leads to differential immune responses in the mesophyll**

**a**, Schematic workflow for identifying *atg5*-specific immune genes. Genes enriched in the immune cluster M11 versus other mesophyll cells were extracted from Col-0_*Pst*_2dpi and pre-activated atg5_mock_2dpi samples. Overlapping genes were subjected to GO enrichment, and immunity-related clusters were visualized as a heatmap and selected as reporter candidates.

**b**, GO enrichment for overlapping genes from (a)
**c,e,g,h** Confocal microscopy images of 5-week-old *A. thaliana* wild type and *atg5* plants expressing *pPEN3::nls-3xGFP* (c), *pSDF2::nls-3xGFP* (e), *pPR2::nls-3xGFP* (g), *pADR1_L1::nls-3xGFP* (h) syringe-infiltrated with 5 x $10^7$ CFUs/mL *Pst ΔhrcC* or mock. Leaves were harvested at 1dpi and fluorescent images are maximal intensity projections of Z-stacks. Scale bar represents 100 µm.
**d**,**f** quantification for (c) and (e), respectively. Different letters indicate statistically significant differences, determined by two-way ANOVA followed by Tukey HSD post-hoc tests (adjusted $p < 0.05$). This experiment was repeated three times with similar results.
**j**, **k**, Confocal microscopy pictures of plants expressing 5-week-old *A. thaliana pSDF2::nls-3xGFP* (j), *pEDS1::nls-3xGFP* (k). Left panel, show the expression in epidermis and guard cells. Right panel, show the expression from mesophyll and vasculature cells. j, star point the stomata positions. Scale bars indicate 100 µm
**i**, heat map showing the relative expression of up_by_*Pst* genes from (a) across cell types in all mock_Col-0 samples.

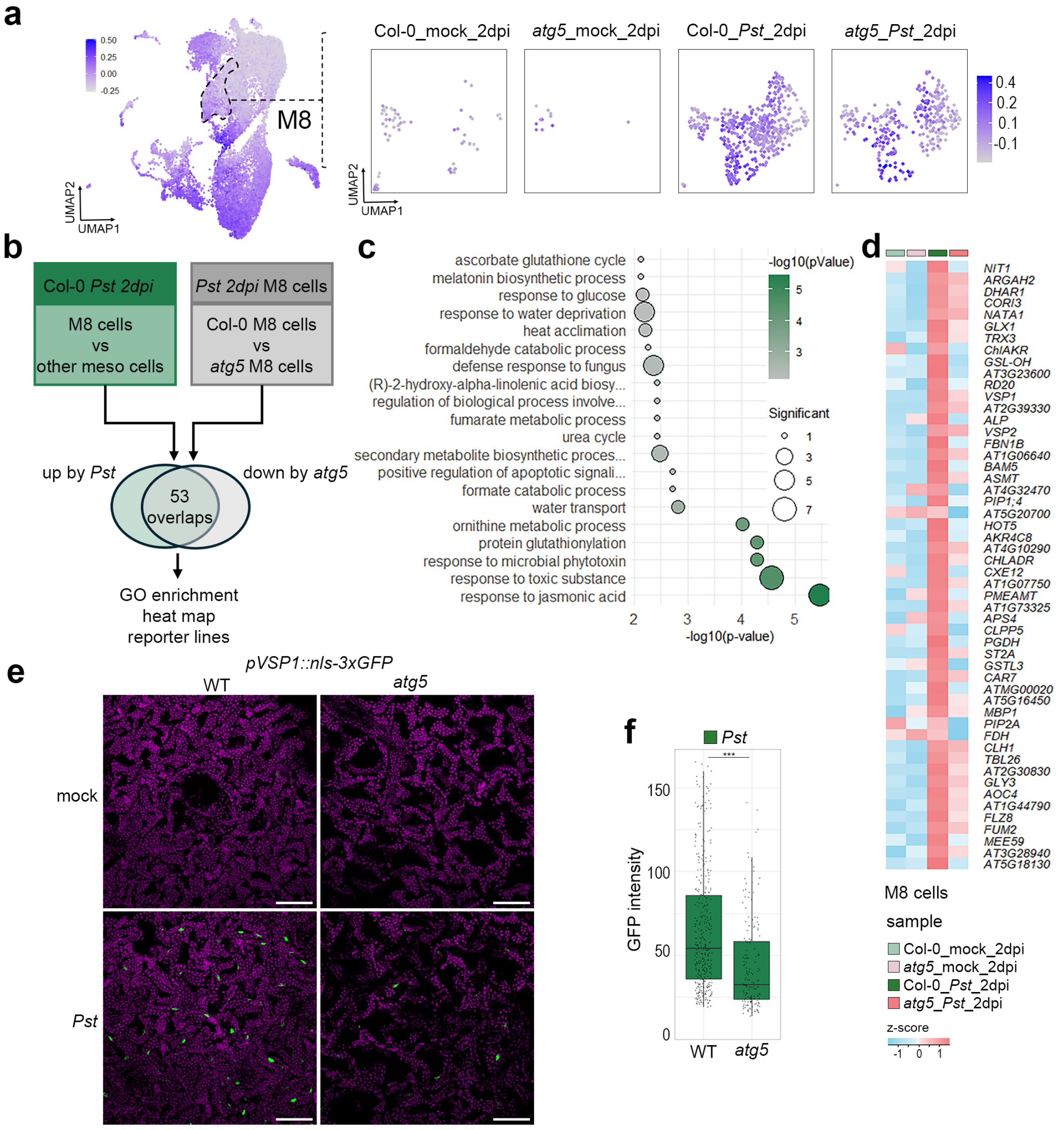


**Extended Data Fig. 6: A disease-associated mesophyll transcriptional program is attenuated in autophagy-deficient plants**

**a**, UMAP plot visualizing Pst_responsive score across M8 cluster cells. Left panel: dashed line outlines the M8 disease-related clusters. Right panel: UMAP for the four 2dpi samples: Col-0 _mock _2 dpi, *atg5*_mock_2 dpi, Col-0_*Pst*_2 dpi, and *atg5*_*Pst*_2 dpi .
**b**, Schematic workflow for the selection and analysis of *atg5*-disease-specific genes. Genes that are significantly enriched in disease cluster M8 compared with other mesophyll cells were obtained for infected sample Col-0_*Pst*_2dpi. The genes that are significantly highly expressed in M8 Col-0_*Pst*_2dpi that that of atg5_*Pst*_2dpi sample were obtained. GO enrichment analysis were followed for the overlap genes from the two obtained datasets and visualized in heatmap and selected for reporter lines candidates.
**c**, GO enrichment for overlapping genes from (b).
**d**, heat map showing the relative expression for overlapping genes from (b) in M8 cluster of the same four samples.
**e**, Confocal microscopy pictures of 5-week-old *A. thaliana* wild type and *atg5* plants expressing *pVSP1::nls-3xGFP* dip-inoculated with 2x10$^{7}$ CFUs/mL *Pst* or 10 mM $MgCl_2$ mock at 4 dpi.
**f**, quantification for (e). Significant differences between genotypes are accessed withStudent's t-test (***$p \leq 0.001$).

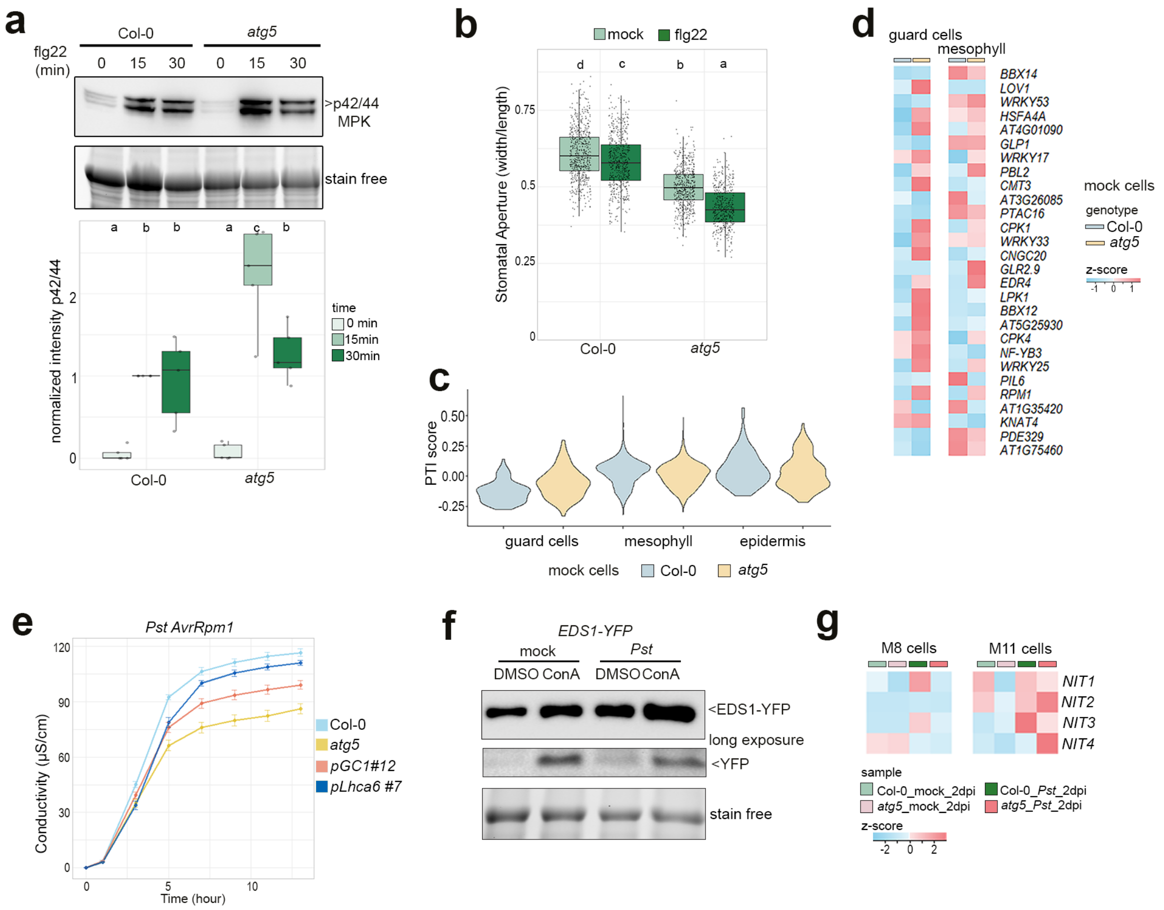


**Extended Data Fig. 7: Autophagy promotes immune potentiation in mesophyll cells**

**a**, immunoblot analyses of MPK activation. 12-day-old Col-0 and *atg5 A. thaliana* seedlings were treated with 10 µM flg22, and samples were collected 0 to 30 min after treatment as indicated. Top panel, activated MAPKs were detected by immunoblotting using anti-p44/42 MAPK antibody. Stain-Free imaging serves as a loading control. Lower part: Quantification of phosphorylated MPKs band intensity from 5 independent experiments.
**b**, Stomatal apertures of 5-week-old *A. thaliana* Col-0, and *atg5* plants after treatment with 5 µM flg22. a.b, different letters indicate statistically significant differences in each genotype, determined by two-way ANOVA followed by Tukey HSD post-hoc tests (adjusted $p < 0.05$).

**c**, Violin plots showing the distribution of the PTI-score across in indicated cell-types and genotypes. The shape illustrates the density of the score distribution for each sample.

**d**, Heat map showing relative expression of PTI induced genes in indicated cell-types and genotypes.

e, Electrolyte leakage over time in 5-week-old *A. thaliana* Col-0, *atg5*, *atg5*; *pLhca6::ATG5-GFP* and *atg5*; *pGC1:: ATG5-GFP* plants following syringe-inoculation with 2 x$10^7$ CFUs/mL *Pst* expressing AvrRpm1. Lines represent means and error bars indicate standard error of the mean from 6 biological replicates of 6 leaf discs each. Similar results were obtained in two independent experiments.

f, EDS1 degradation assay revealed by the ratio of free YFP to full-length EDS1-YFP. Immunoblot against GFP on crude extracts of 10-day-old *A. thaliana* seedlings EDS1-YFP in *eds1-2* background, infected with *Pst* or 10 mM $MgCl_2$ mock for 8 hours following overnight 1 µm concanamycin A. GFP blot was split for visualization purposes

**g**, Heat map showing the relative expression of NIT genes in M11 and 8 cluster of the same four samples.
